# Nav1.3 and fibroblast growth factor homologous factor 14 are primary determinants of the TTX-sensitive sodium current in mouse adrenal chromaffin cells

**DOI:** 10.1101/2020.10.31.363416

**Authors:** P.L. Martinez-Espinosa, C. Yang, X.M. Xia, C.J. Lingle

## Abstract

Adrenal chromaffin cells (CCs) in rodents express a rapidly inactivating, TTX-sensitive sodium current. The current has generally been attributed to Nav1.7, although a possible role for Nav1.3 has also been suggested. Nav channels in rat CCs rapidly inactivate into two separable pathways, which differ in their time course of recovery from inactivation. One population recovers with time constants similar to traditional fast inactivation and the other about 10-fold slower. Inactivation properties suggest that the two pathways result from a single homogenous population of channels. Here we probe the properties and molecular components of the Nav current present in mouse CCs. We first confirm that functional properties of Nav current in rat and mouse cells are generally similar in terms of activation range, steady-state inactivation, and dual pathway fast inactivation. The results then show that all inward Nav current is absent in CCs from Nav1.3 KO mice. Subsequently, in a mouse with KO of fibroblast growth factor homology factor 14 (FGF14), we find that the slow component of recovery from fast inactivation is completely absent in most CCs, with no change in the time constant of fast recovery. Experiments probing the use-dependence of Nav current diminution between WT and FGF14 KO mice directly demonstrate a role of slow recovery from inactivation in determination of Nav current availability. Overall, the results indicate that the FGF14-mediated inactivation is the major determinant in defining use-dependent changes in Nav availability in CCs. We also consider the potential impact that inactivating FGF’s with different recovery kinetics can exert on differential use-dependent changes in Nav availability.

## Introduction

Adrenal chromaffin cells express a rapidly inactivating TTX-sensitive Nav current (Fenwick et al., 1982; Islas-Suarez et al., 1994; Vandael et al., 2015). Despite the rapid inactivation of CC Nav current and rapidly activated repolarizing K^+^ currents (Martinez-Espinosa et al., 2014; Lingle et al., 2017), depolarization-evoked action potential (AP) firing frequency in CCs is typically limited to about 10-20 Hz (Solaro et al., 1995; Martinez-Espinosa et al., 2014). In an associated paper (Martinez-Espinosa et al., 2020), following brief 5 ms inactivation steps, Nav current in rat CCs was shown to recover from inactivation with two separable processes, one with a fast recovery of ~3-30 ms and the other ~50-300 ms, each dependent on voltage. This dual fast inactivation process is similar to that which has been termed “long-term inactivation” (Goldfarb, 2012; Barbosa and Cummins, 2016), for which the slow recovery component has been proposed to arise from inactivation involving N-termini of particular isoforms of intracellular fibroblast growth factor homologous factors (FHFs or FGFs) (Dover et al., 2010). The fast inactivation process leading to slow recovery is thought to occur in a largely competitive fashion with typical intrinsic fast inactivation. The slow recovery process provides a potential mechanism by which Nav channel availability may be reduced during repetitive activity, perhaps influencing the firing frequencies observed in CCs (Venkatesan et al., 2014; Navarro et al., 2020). In order to better understand the relationship between intrinsic fast inactivation and any competing fast inactivation process, it is important that the underlying molecular entities be defined. Here we address the question of the molecular identity of the Nav channel pore-forming subunits in rodent CCs and also evaluate the role of FGF14 in the inactivation process.

The initial cloning of a neuroendocrine sodium channel, both from human and rat tissues, (Klugbauer et al., 1995) established the presence of Nav1.7 (*Scn9a*) message in adrenal chromaffin cells. Subsequent work has supported this idea, demonstrating the presence of Nav1.7 message and protein, primarily in dissociated bovine CCs (Wada et al., 2004; Nemoto et al., 2013; Tamura et al., 2014), and has also identified pathways that can up- or down-regulate Nav1.7 expression (Wada et al., 2008). However, a recent paper using quantitative RT-PCR found that Nav1.3 message was more abundant than Nav1.7 in mouse CCs (Vandael et al., 2015). Furthermore, putative protein for each was detected in western blots from adrenal medulla (AM) (Vandael et al., 2015), although samples from KO animals was not evaluated.

When both Nav1.3 and Nav1.7 currents have been found in the same cells, as in mouse pancreatic alpha and beta cells, steady-state inactivation curves reveal two distinct Boltzman components that are attributable to separate contributions of Nav1.7 and Nav1.3 currents, as confirmed with Nav1.3 and Nav1.7 KO mice (Zhang et al., 2014). In contrast, Nav currents in CCs, whether mouse or rat, appear to have steady-state inactivation properties more consistent with only a single component, more similar to Nav1.3 (Lou et al., 2003; Vandael et al., 2015; Martinez-Espinosa et al., 2020). To date, heterologous expression studies have not been particularly useful regarding to the extent to which Nav1.3 and Nav1.7 can be readily distinguished. Heterologously expressed Nav1.3 and Nav1.7 have not been studied side-by-side in the same study, but when Nav1.3 (Cummins et al., 2001) and Nav1.7 (Cummins et al., 1998; Herzog et al., 2003; Cummins et al., 2004) have been expressed in HEK cells, the steady-state inactivation curve for Nav1.7 is left shifted (V_h_ ~−75 mV) compared to Nav1.3 (V_h_~−65 mV). Whether a difference between Nav1.7 and Nav1.3 components is readily resolvable is not clear. The utility of such comparisons can also be affected by what Nav β subunits may be present in native cells or employed in heterologous expression studies.

Motivated by the potential role of dual-pathway fast inactivation and the slow component of recovery from inactivation in regulation of AP firing in rat CCs (Martinez-Espinosa et al., 2020), here we have turned to mouse CCs to begin to tease apart the molecular components of Nav current. We first compare functional properties of Nav current in mouse and rat CCs. We then show that all Nav current in mouse CCs is abolished in the Nav1.3 KO mouse. Furthermore, using FGF14 KO mice, we examined whether an FGF14 isoform might be responsible for the slow component of recovery from inactivation in rodent CCs. We find that, in FGF14 KO mice, the slow component of recovery from inactivation is absent (about 75% of tested cells) or markedly reduced. Given earlier results establishing a role for FGF14A, but not FGF14B, in use-dependent inactivation and kinetics of slow recovery from inactivation (Laezza et al., 2009; Dover et al., 2010), most likely FGF14A is the primary determinant of dual-pathway fast inactivation in mouse CCs. Overall, the results indicate that Nav1.3 and FGF14 together are the primary molecular components that underlie Nav channel inactivation behavior in mouse CCs, although an additional FGF subunit may contribute to Nav channels in some cells. The results also demonstrate directly that the presence of FGF14 impacts substantially on changes in Nav channel availability during repetitive stimuli.

## Methods

### Animals

Mice (8-12 week old) were sacrificed by CO_2_ inhalation, following protocols approved by the Washingon University in St. Louis Institutional Care and Use Committee. Animals were housed in accordance with the National Institutes of Health Committee on Laboratory Animal Resources guidelines. Nav1.3 KO mice (C57BL6j background) were kindly provided by Dr. Steven Waxman and D.-H. Sulayman (Yale University) and maintained as homozygous knockout mice. Control C57BL6 mice were obtained from Jackson Labs. FGF14 KO mice (Wang et al., 2002) were kindly provided by Dr. Jeanne Nerbonne with permission of Dr. David Ornitz (both of Washington University Sch. Med.). For FGF14 KO mice in which a slow component of recovery from inactivation was noted, mice were genotyped multiple times by two individuals (one in the Lingle lab and one in the Nerbonne lab) using different sets of primers. In all cases, regenotyping of the putative FGF14 KO mice confirmed the original genotype assignment.

### Cell preparations

All experiments were done on CCs in adrenal medullary slices with slice preparation and solutions as described in the associated paper (Martinez-Espinosa et al., 2020).

### Electrophysiological techniques

Whole-cell recordings from adrenal slices were done within 5 hours after isolation of the adrenal gland. For recordings of Nav current, the standard open-pipette method (Hamill et al., 1981) method was employed. Whole-cell voltage-clamp recordings were accomplished with a Multiclamp 700B amplifier (Molecular Devices, Sunnyvale, CA). Command waveforms and data acquisition was done with the Clampex program from the pCLAMP 9.0 software package (Molecular Devices). Currents were evaluated without leak subtraction. Time constants of exponential relaxations in current records were fit using Clampfit algorithms. Fitting of Boltzmann functions or exponential recovery time course was done either using Excel or with a custom program using a Levenberg-Marquardt algorithm for non-linear least-squares fitting. Normalized GV curves were generated from: G(V)=I/(V_m_-V_r_) with V_r_=66 mV, which assumes linearity in instantaneous current over voltages up to about +10 mV.

Typical membrane capacitance for mouse chromaffin cells included in this study was 8.95 ± 0.48 pF. Patch-clamp micropipettes were pulled from borosilicate glass (Drummond). Pipette resistances ranged from 1.5-2.5 MΩ. Electrodes were coated with Sylgard 184 (Dow Chemical) and fire polished. The reference electrode was a Ag/AgCl_2_ pellet in direct contact with the bath.

### Recording solutions

The internal saline had the following composition (in millimolar): 125 CsCl, 10 NaCl, 5 EGTA, 4 Mg-ATP and 10 HEPES (pH 7.4 with CsOH) The standard extracellular solution contained (mM) 119 NaCl, 23 NaHCO_3_, 1.25 NaH_2_PO_4_, 5.4 KCl, 2.0 MgSO_4_, 1.8 CaCl_2_, 11 glucose, 2 sodium pyruvate, and 0.5 ascorbic acid, pH 7.4. Membrane capacitance (C_m_) and series resistance (R_s_) were read from amplifier settings and R_s_ compensation set to 90%. Analysis of currents have been limited to cells in which the estimated maximal voltage error resulting from residual uncompensated R_s_ was less than 10 mV (Martinez-Espinosa et al., 2020) Perfusion of external salines was performed by switching the solution flowing into the slice chamber.

### RNA Extraction and Quantitative RT-PCR

For quantitative PCR measurements, four Sprague-Dawley rats (300-324 grams, about 70-75 days old) were ordered from Harlan Laboratories. Mice were C57/BL6 (8-12 weeks old). Following sacrifice by CO_2_ inhalation, adrenal glands were dissected under a microscope. The cortex was carefully removed and the AM was quickly frozen in liquid nitrogen and then saved at −80°C. Total RNA from each pair of AM from a single animal was isolated using the RNeasy Plus Mini Kit (Qiagen). cDNA was synthesized using the BioRad iScript cDNA Synthesis Kit (Cat. No. 170-8891). For the negative control groups, all components except the reverse transcriptase were included in the reaction mixtures. Real-Time PCR was performed with specific primers purchased from Qiagen (**Tables 1,2**) and Power SYBR Green PCR Master Mix (Applied Biosystems, Cat. No. 4367659) under reaction conditions identical to those described previously (Yang et al., 2009). PCR specificity was verified by a dissociation curve with a single peak which was run following the real-time PCR reaction. Message levels were normalized to the abundance of β-actin message. The mean value was averaged from at least 3 separately prepared RNA samples, with each sample run in triplicate.

**Table 1.** Primers used for real-time PCR in rat tissues.

| rat gene | Cat.No. of QuantiTect Primer Assay from Qiagen | amplicon length | amplified exons |
| --- | --- | --- | --- |
| *scn1a* | QT02445618 | 134 bp | 20/21 |
| *scn2a* | QT00190722 | 66 bp | 14/15 |
| *scn3a* | QT01568126 | 86 bp | <sup>3</sup>/<sub>4</sub> |
| *scn4a* | QT00184786 | 113 bp | 12/13 |
| *scn8a* | QT00195496 | 67 bp | 7/8 |
| *scn9a* | QT00194558 | 101 bp | 16/17 |
| *scn1b* | QT00181559 | 96 bp | $\frac{3}{4}$ |
| *scn2b* | QT00193417 | 140 bp | $\frac{1}{2}$ |
| *scn3b* | QT00179459 | 118 bp | 4/5 |
| *scn4b* | QT01581349 | 93 bp | $\frac{1}{2}$ |
| *gapdh* | QT00199633 | 149 bp | 1/3 |
| *actB* | QT00193473 | 145 bp | 2/3 |

**Table 2.**
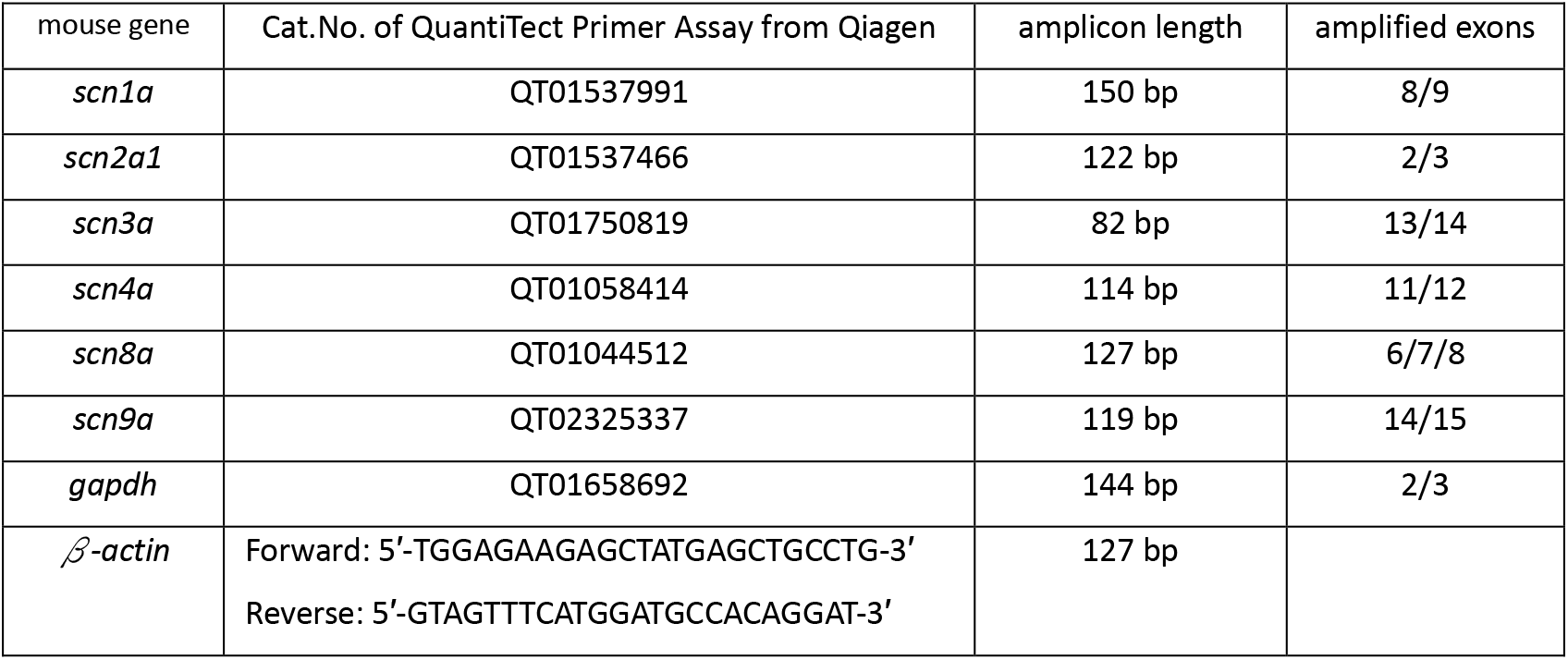
Primers used for real-time PCR in mouse tissues.

### Statistics

Unless otherwise indicated in a figure legend, most graphs display means ± SD for sets of measurements obtained from different cells from a given genotype. All reported N values refer to numbers of cells, although in some cases numbers of animals from which the cells were taken is also reported. Given that our results directly demonstrate that key aspects of the inactivation behavior under investigation involve heterogeneity among cells arising from unknown molecular components of the channels, grouping of all cells from a single animal into an averaged measurement will artifactually lead to misinterpretation about the basic underlying phenomena. Thus, in regards to evaluating the behavior of a given type of current among cells and animals from a given genotype, a cell appears to be the appropriate experimental unit for consideration. In some cases, functions (Gaussian, exponential, Boltzmann) were used to fit curves. In such cases, best fit parameters are reported along with the 90% confidence limit on the fitted parameter. For statistical tests, if more than 10 values are available for a comparison between two distributions, we employ a Kolgoromov-Smirov test (KD-test), since this test makes no assumptions about normality. In other cases, we employ an ordinary ANOVA with Tukey’s corrections for multiple comparisons (Graphpad Prism 8, San Diego CA). Exact P values are reported in the Legends, except when P<0.001. Readers will notice that different figures report results for recovery from inactivation at −80 mV with different numbers of WT cells. As is typical, not all protocols were tested on all cells. As such, in comparisons of voltage-dependence of recovery from inactivation (Figs. 2–3), we only included cells for which recovery was examined at more than one recovery voltage, resulting in a set of 16 WT cells for which recovery at −80 mV was obtained along with at least one other recovery voltage (−60, −100, or −120 mV). However, when comparing recovery between WT to FGF14 KO cells (Fig. 6–7), we included all WT cells from which recovery was examined at −80 mV (a total of 33 WT cells), irrespective of whether recovery was examined at other voltages. Finally, Fig. 9 only includes cells (for WT, N=22) in which recovery at −80 mV was measured both with the 1P and 10P protocols.

## Results

### Properties of Nav currents in mouse CCs recorded in adrenal medullary slices

Figure 1A shows Nav current activated in a CC from a mouse adrenal medullary slice, with a 5 ms activation step over voltages from −70 to +50 mV, following a 1000 ms conditioning step to −120 mV to remove any channel inactivation. Average peak Nav current density for mouse cells (N=18) was determined and compared to similar results for rat cells (Fig. 1B), with average Nav current density in mouse CCs about 70% of that in rat CCs. Normalized current amplitude (Fig. 1C) shows that both mouse and rat Nav currents activate over a similar range of voltages, although the mouse currents are somewhat right shifted compared to rat cells. Conversion of the current-voltage relationship to conductance-voltage (GV) curves, assuming a +66 mV reversal potential, resulted in GV curves that were similar, but not identical (Fig. 1D). For mouse, the voltage of half activation (V_h_) was −22.3±0.3 mV with *z*=5.3±0.3*e* and, in rat, V_h_=−27.4±0.2 mV with *z*=5.4±0.2*e*. Although such GV curves based on peak currents distort the true GV properties of the current because of the effects of inactivation on the peak current, this apparent mouse GV is shifted about +5 mV relative to rat (KS test P =0.004). The rate of onset of inactivation, measured from single exponential fits to the decay phase of the Nav current, was generally similar to that in rat (Fig. 1E). Comparison of the steady-state inactivation curves generated with 25, 100, 250 and 1000 ms conditioning duration over voltages from −100 through 0 mV exhibited a leftward shift with conditioning pulse duration qualitiatively similar to that observed in rat chromaffin cells (Martinez-Espinosa et al., 2020). However, over conditioning durations of 100-1000 ms, the mouse fractional availability curve was shifted compared to rat cells about −5 to −9 mV (Fig. 1G-H). With a 1000 ms conditioning duration, for mouse, V_h_=−48.9±1.0 mV, while, for rat, V_h_=−57.7±0.6 mV. Some of this presumably reflects coupling to the shifted activation range. Similar to rat cells, there was little indication that a double Boltzman function would better describe the steady-state inactivation behavior, except for a small indication of a more negative component with the 25 ms conditioning step. It is important to recognize that peak current elicited from a conditioning potential of −80 mV and that from −120 mV differed by less than 5% in both rat and mouse. This is markedly different from the behavior of mouse pancreatic β and α cells which have been unambiguously shown to express both Nav1.3 and Nav1.7 currents (Zhang et al., 2014)

**Figure 1.**
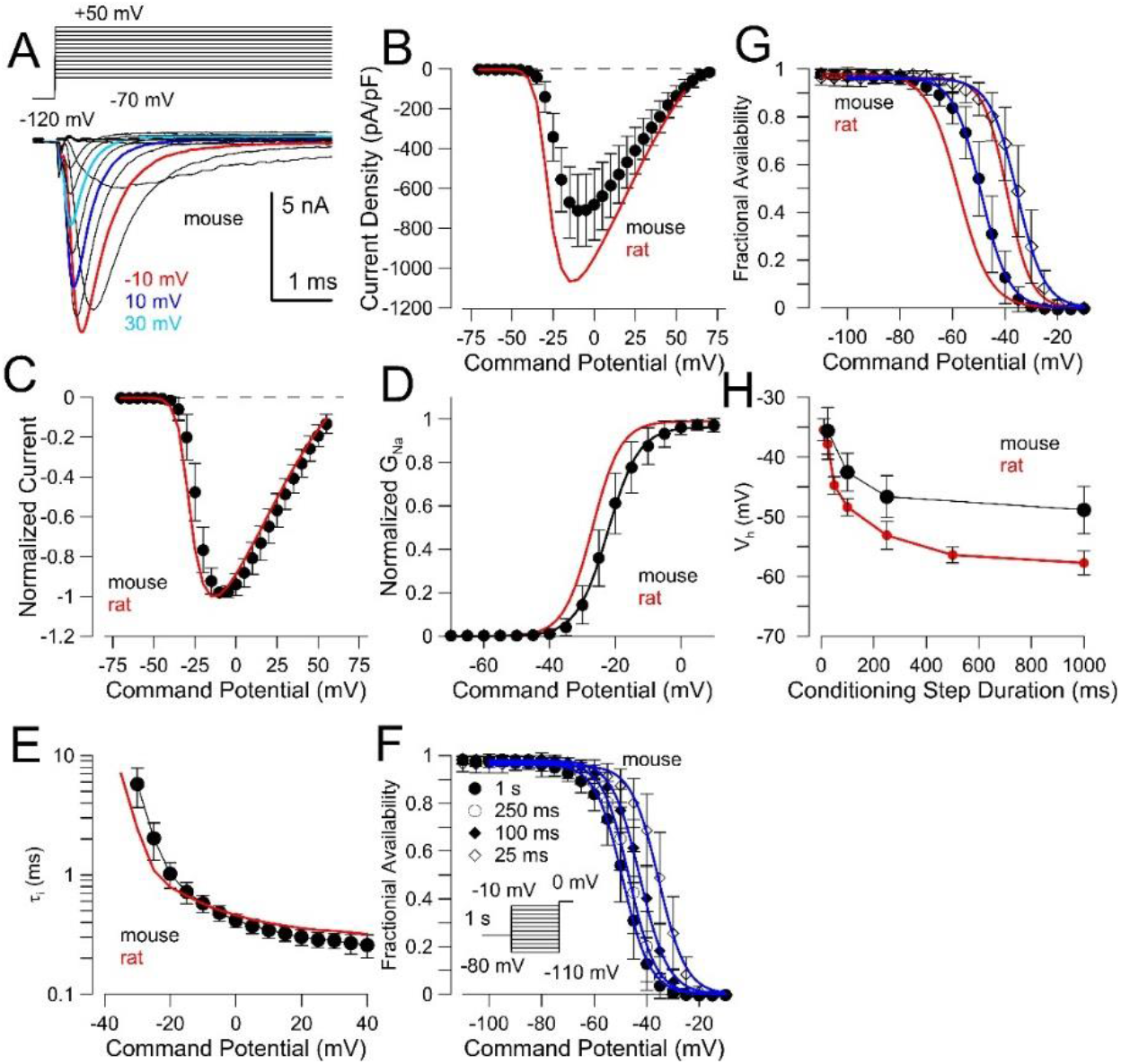
Basic properties of Nav current in mouse CCs are similar to those in rat CCs. (A) Example currents illustrating basic voltage-dependent activation in a mouse CC in a slice. (B) Plot of Nav current density (mean±SD) from 18 mouse CCs. Red line corresponds to current density from rat CCs. (C) Points show normalized peak current amplitude for mouse CCs, compared to rat CCs (red line). (D) Mouse (and rat) peak currents were converted to conductances assuming a reversal potential of +66 mV. For the GV from mouse CCs, V_h_=−22.3±0.3 mV with *z*=5.3±0.3*e*, while, for rat, V_h_=−27.4±0.2 mV with *z*=5.4±0.2*e.* (E) Inactivation time constants (mean±SD) as a function of command potential are plotted for 18 mouse CCs and compared to values from rat (red line). (F) Steady-state inactivation curves following 25, 100, 250, and 1000 ms conditioning steps at voltages from −110 to 0 mV for mouse. For 25 ms, V_h_ = −35.5±0.3 mV, z=4.8±0.3*e;* for 100 ms, V_h_=-42.6±0.3 mV, z=4.7±0.2*e*; for 250 ms, V_h_=−46.7±0.2 mV, 5.0±0.2*e*; for 1 s, V_h_=−49.2±0.3 mV, z=4.8±0.2*e*. (G) Comparison of mouse and rat (red) steady-state inactivation curves following 25 ms and 1000 ms conditioning steps. At 25 ms for rat cells, V_h_=−37.8 ± 1.3 mV and, at 1000 ms, V_h_=−57.7±0.6 mV. (H) Comparison of V_h_ of fractional availability following conditioning steps of differing durations for mouse and rat (red).

### Comparison of fast and slower recovery from inactivation in rat and mouse CCs

The distinguishing feature of Nav current in rat CCs is a dual-pathway fast inactivation process that leads to two distinct components of recovery from inactivation (Martinez-Espinosa et al., 2020). Here we examined recovery from inactivation in mouse CCs. Using a standard paired pulse recovery protocol, Figure 2 illustrates the time course of recovery from inactivation in a mouse CC for recovery at −80 (Fig. 2A) and −120 mV (Fig. 2B). The time course of recovery from inactivation in the mouse CCs was well-described by a double exponential recovery process (Fig. 2C) comparable to that in rat CCs (Fig. 2D), albeit with some differences in absolute time constants.

**Figure 2.**
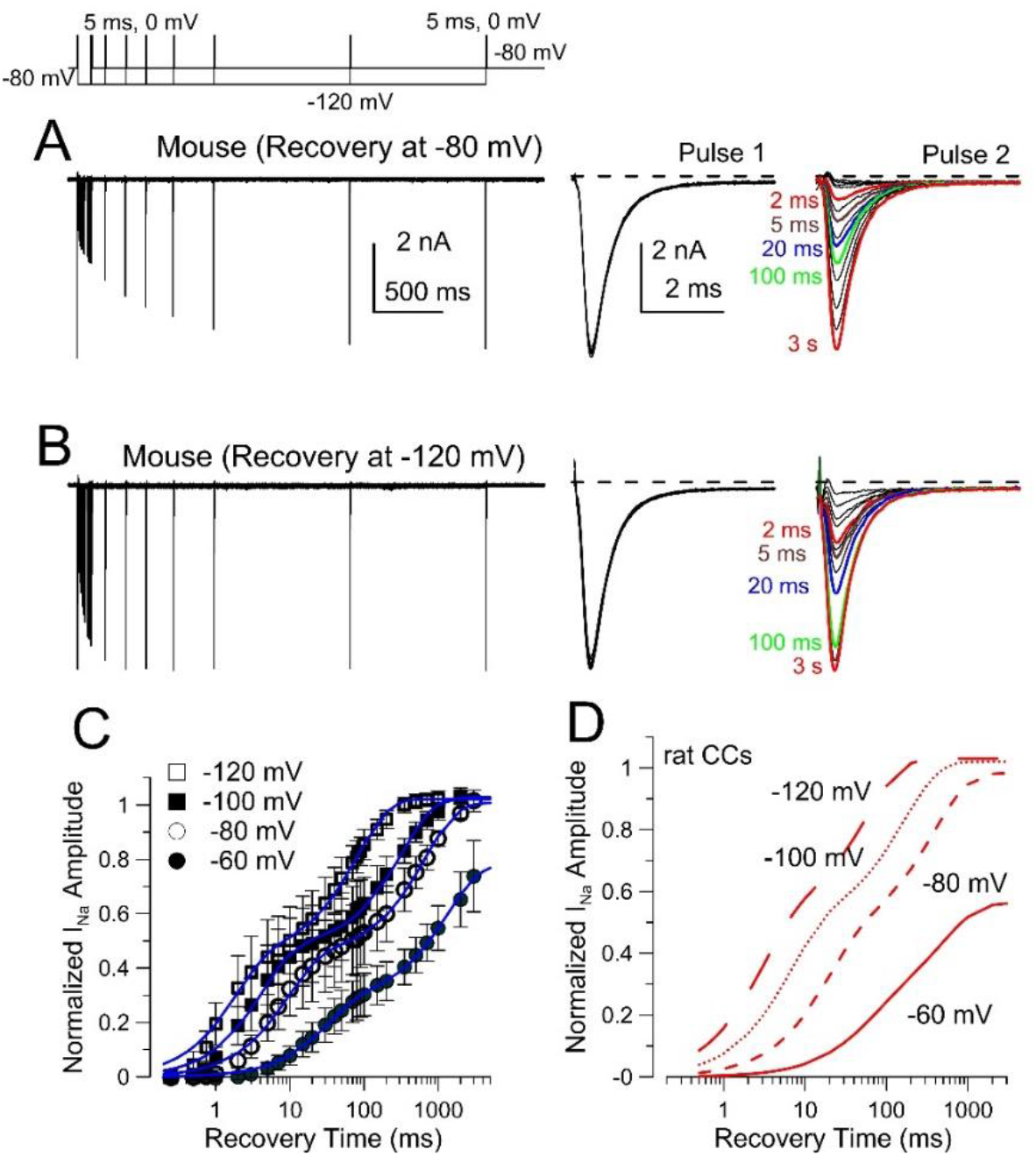
Nav current in mouse CCs exhibits two components of recovery from inactivation, generally similar to that in rat CCs. (A) Example paired pulse (PP) recovery protocol with recovery at −80 mV following a 5 ms inactivation step at 0 mV. On the right, aligned P1 and P2 currents are shown. (B) Traces show the time course of recovery from inactivation at a −120 mV recovery potential for the same cell as in A. (C) Averaged time course (±SD) of recovery for mouse CCs at the indicated voltages along with best fit double exponential function. For −120 mV (12 cells), A_f_=0.46±0.02, t_f_=2.17±0.31 ms, A_s_=0.56±0.02, t_s_=83.3±10.2 ms. For −100 mV (n=12 cells), A_f_=0.48±0.02, t_f_=4.38±0.49 ms, A_s_=0.54±0.02, t_s_=335.3±43.1 ms. For −80 mV (n=18 cells), A_f_=0.45±0.01, t_f_=9.38±0.75 ms, A_s_=0.56±0.02, t_s_=660.5±70.4 ms. For −60 mV (n=8 cells), A_f_=0.29±0.01, t_f_=32.4±2.7 ms, A_s_=0.49±0.02, t_s_=1336.4±176.0 ms. (D) Fits of recovery from inactivation in rat CCs from associated paper (Martinez-Espinosa et al., 2020).

Variation in the time course of recovery from inactivation at −80 mV (Fig. 3A) and −120 mV (Fig. 3B) was compared more closely among individual cells. The average fraction of the fast recovery component was near 0.5 at either −80 mV or −120 mV (Fig. 3C) comparable to that in rat cells. For an individual cell, the amplitudes of fast recovery measured at different recovery voltages was similar (dotted lines in Fig. 3C). Overall, the fractional amplitude of the fast recovery component (Fig. 3C) and mean time constants (Fig. 3D-F) were similar between species, although slow recovery from inactivation at −80 mV is slower in mouse than in rat (P<0.0001; Fig. 3E). Overall, the key features of the two component recovery are generally comparable between mouse and rat, perhaps consistent with the idea that they have similar molecular underpinnings.

**Figure 3.**
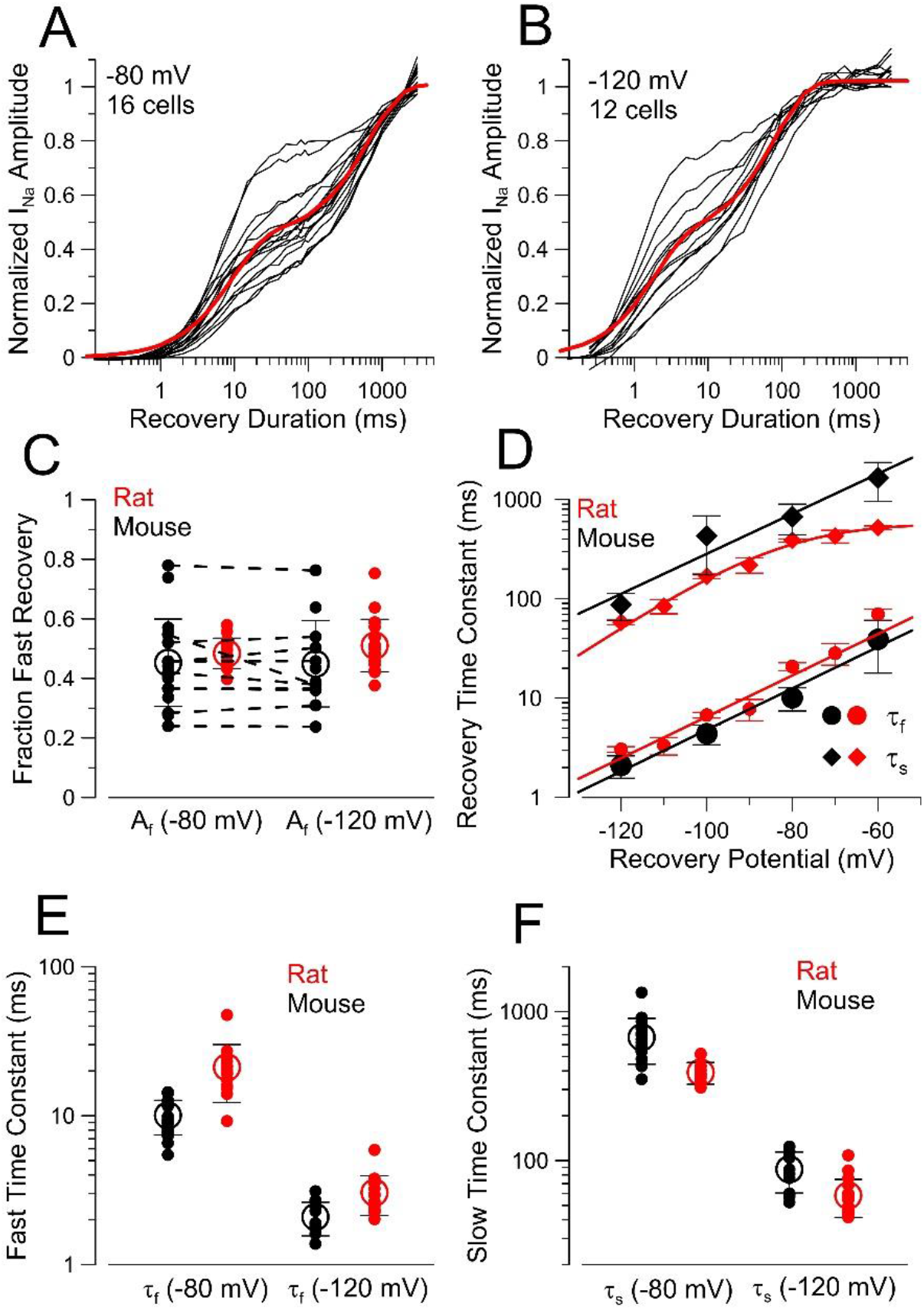
Comparison of two component recovery from inactivation in mouse and rat. (A) The time course of recovery from inactivation at −80 mV for 16 WT mouse CCs is shown, along with the averaged recovery time course (red line, from Fig. 2C). (B) Recovery from inactivation at −120 mV for 12 WT CCs, along with averaged recovery (red line). (C) The fraction of the fast recovery component at −80 mV or −120 mV is plotted for both individual mouse (black) and rat (red) CCs. Dotted lines indicate values obtained in individual cells where both −80 mV and −120 mV recoveries were obtained. Statistical comparisons of distributions of fast recovery amplitude at −80 or −120 mV for both mouse and rat revealed no differences. (D) Mean values (±SEM) for fast (t_f_) and slow (t_s_) recovery time constants are plotted as a function of recovery potential for both mouse and rat CCs. Lines are derived from exponential fits, but only intended to highlight trends in the data. (E) Comparison of tf at −80 mV and −120 mV between mouse and rat CCs, with small symbols corresponding to values from individual cells (F) t_s_ at −80 mV and −120 mV for mouse and rat CCs. Using ANOVA with a Bonferroni correction for multiple comparisons yielded P<0.0001 for comparison of mouse and rat slow time constants at −80 mV, but all other comparisons (fast time constants, fraction fast amplitude, and slow time constants at −120 mV) were P>0.85.

### Evaluation of message for Nav subunits in mouse and rat AM

We next turned to an evaluation of message for Nav channel subunits in rat and mouse adrenal medulla. A previous paper that examined message for Nav variants in mouse AM revealed a more than 5-fold higher level of message for Nav1.3 (*Scn3a*) than Nav1.7 (*Scn9a*) and also reported the presence of bands on western blots perhaps corresponding to both Nav1.3 and Nav1.7 (Vandael et al., 2015). Here, we also tested for the presence of message in isolated mouse (Fig. 4A) and rat (Fig. 4B) AM for known TTX-sensitive Nav variants, including Nav1.1 (*Scn1a*), Nav1.2 (*Scn2a*), Nav1.3 (*Scn3a*), Nav1.4 (*Scn4a*), Nav1.6 (*Scn8a*), and Nav1.7 (*Scn9a*). In mouse AM, Nav1.3 message was predominant (Fig. 4A), while Nav1.7 message was not clearly greater than other weakly expressed forms (*Scn1a, Scn2a, Scn4a, Scn8a*). The levels observed here in mouse are similar to what was observed previously (Vandael et al., 2015), with Nav1.3 message being 5-10 fold higher than Nav1.7 message. In rat, Nav1.3 (*Scn3a*) and Nav1.7 (*Scn9a*) were the most abundant of all candidate TTX-sensitive Nav variants (Fig. 4B), with Nav1.7 message levels exceeding that of Nav1.3.

**Figure 4.**
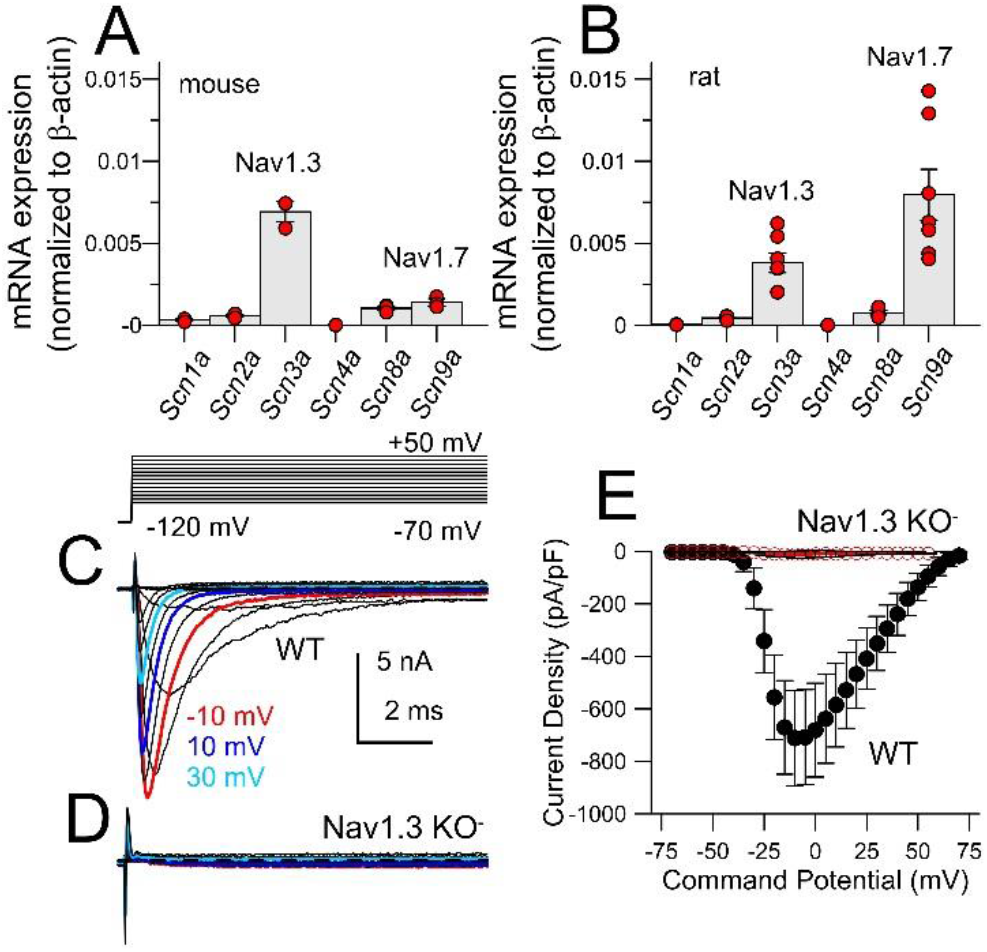
KO of 1.3 (SCN3A) removes all detectable inward Na± current in mouse CCs. (A) Each point represents q-RT-PCR reactions run in triplicate for a single adrenal medulla for the indicated Nav channel subunits from mouse adrenal medulla, with message levels normalized to b-actin. (B) Similar reactions were run on ratadrenal medulla. In both A and B, at least three adrenal medulla were examined for each message species. In A, P values (from ANOVA with Tukey’s posthoc test) of less than 0.05 were: <0.0001 for *Scn2a* vs. *Scn3a*, *Scn3a* vs. *Scn4a*, *Scn3a* vs. *Scn8a*, *Scn3a* vs. *Scn9a*, and 0.0104 for *Scn4a* vs. *Scn9a*. In B, adjusted P values of less than 0.05 were: <0.0001 for *Scn1a* vs. *Scn9a*, 0.0002 for *Scn2a* vs. *Scn9a*, 0.0181 for *Scn3a* vs. *Scn9a*, and 0.0003 for *Scn8a* vs. *Scn9a*. (C) Example Nav current activation from a WT mouse CC. (D) Example currents from a CC from an Nav1.3 KO mouse. Currents in C and D were not leak subtracted. (E) Average peak inward current density (±SD) for CCs from WT (18 cells) and Nav1.3KO (10 cells) mice.

### KO of Nav1.3 abolishes Nav current in mouse CCs

Given the abundance of Nav1.3 message in mouse adrenals, we took advantage of Nav1.3 KO mice kindly provided by Drs. S. Waxman and D.-H. Sulayman (Yale University). Figure 4C and 4D compare whole-cell current recordings with the standard activation protocol for a WT CC and a CC from an Nav1.3 KO mouse, revealing a complete absence of rapidly activated, inactivating inward current in the absence of Nav1.3 (Fig. 4D). Similar results were obtained from a set of 10 cells (Fig. 4E). In some cells, a small non-inactivating component of current was observed with the salines used here. This current most likely represents the small amount of Ca^2+^ current present in rodent CCs with 1.8 mM extracellular Ca^2+^, but contributes inconsequentially to comparison of IV curves between CCs from WT and Nav1.3 KO mice. Thus, genetic ablation of Nav1.3 results in the complete absence of TTX-sensitive Nav current in mouse adrenal CCs.

### FGF14 underlies the slow component of recovery from inactivation in mouse CC

Long-term Nav inactivation described in other systems (Dover et al., 2010; Navarro et al., 2020) has been shown or proposed to involve inactivation mediated by intracellular fibroblast growth factor homologous factors (abbreviated either FGFs or FHFs) (Smallwood et al., 1996; Munoz-Sanjuan et al., 2000). This family of proteins consists of a family of four peptides (FGF11-FGF14, also known as FHF3, FHF1, FHF2, or FHF4, respectively) with splice variation at the N-terminus. Although these share a core segment with strong homology to secreted fibroblast growth factors (Smallwood et al., 1996; Munoz-Sanjuan et al., 2000), in general they appear to not be secreted and to interact with Nav channels and potentially other proteins (Pablo and Pitt, 2014).

Given similaries of dual-pathway fast inactivation in rodent CCs to long-term Nav inactivation, we have taken advantage of the availability of FGF14 KO mice (kindly provided by Dr. Jeanne Nerbonne, Washington University) to compare Nav current properties between WT and FGF14 KO CCs. Nav current in CCs from FGF14 KO mice was generally similar to those from WT CCs (Fig. 5A vs. 5B), with comparable current densities (Fig. 5C). The GV for current Nav current activation in the FGF14KO cells was shifted by about +5 mV (Fig. 5D). We also observed that the time course of the onset of inactivation at voltages from −20 mV to +40 mV (Fig. 5E-F) was faster for FGF14 KO CCs (at 0 mV, τ_i_=0.38±0.02 ms) than WT cells (at 0 mV, τ_i_=0.50±0.03 ms). The steady-state inactivation curve for mouse Nav current in the absence of FGF14 was shifted leftward about −10 mV (Fig. 5G-H). This is qualitatively similar to the approximately −13 mV shift observed for granule cell Nav current following double KO of FGF12 and FGF14 (Goldfarb et al., 2007), while the presence of FGF14A produces an ~10 mV positive shift inactivation V_h_, when coexpressed with Nav1.6 (Laezza et al., 2009).

**Figure 5.**
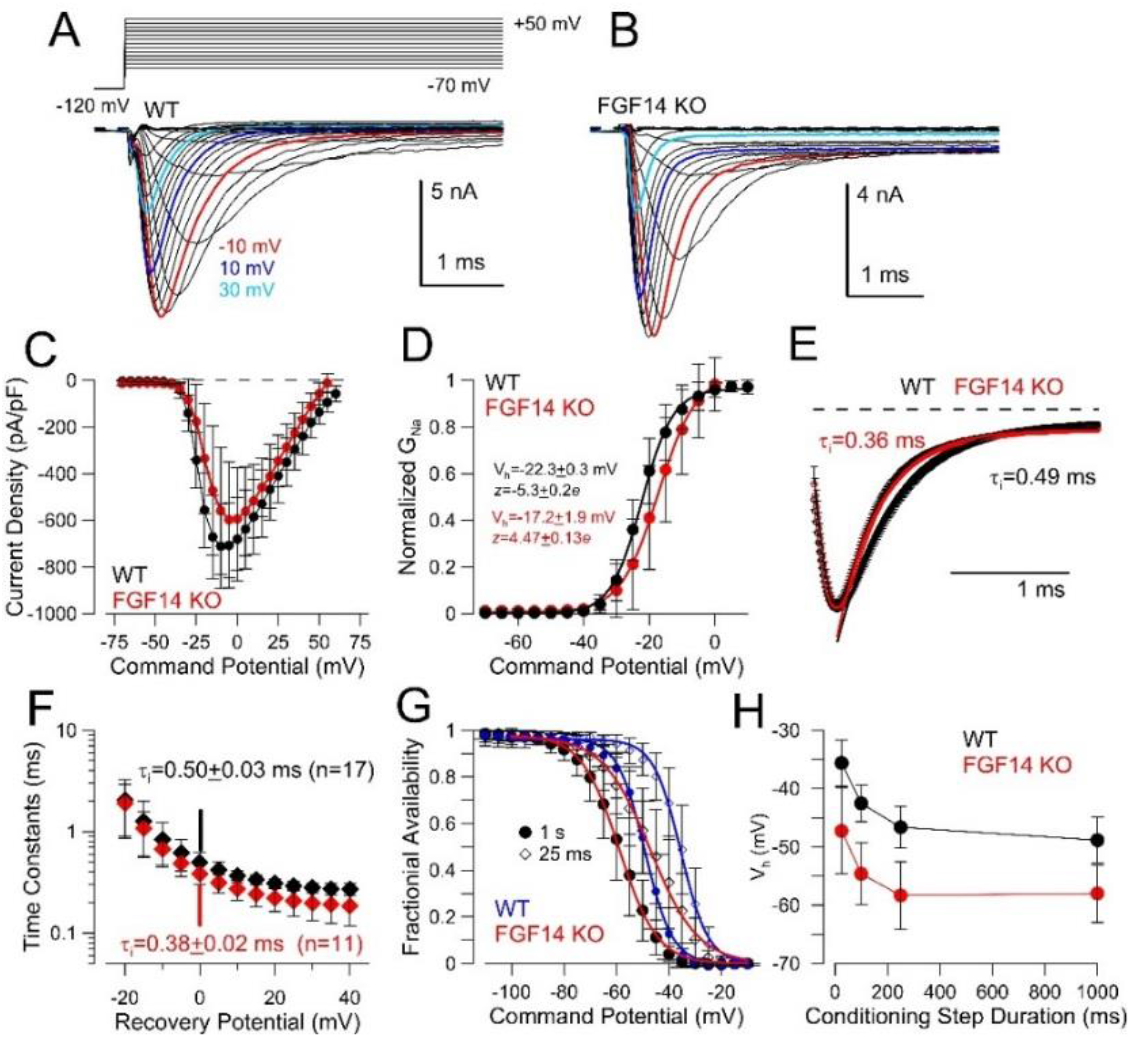
Nav currents in CCs from FGF14 KO animals. **(A)** Currents from a WT mouse CC activated with the indicated protocol with colored traces as indicated. **(B)** Currents from an FGF14 KO CC. **(C)** Voltage-dependence of current density for CCs from FGF14 KO mice (red points; 10 cells) and those from WT mice (18 cells). Mean ± SD. **(D)** Normalized GV curves (see Methods) for WT (from Fig. 1D) and FGF14 KO cells. For WT, V_h_=−22.3±0.3 mV with *z*=5.3±0.3*e* and, for FGF14 KO, V_h_=−17.2±1.9 mV with z=4.5±0.1*e*. **(E)** Traces of Nav current activated at 0 mV from WT (black) and FGF14 KO (red) cells were averaged and fit with single exponential functions, with faster onset of inactivation in FGF14 KO cells. **(F)** Mean values (±SD) for onset of inactivation for WT and FGF14 KO cells are plotted as a function of command potential. P-values adjusted for multiple comparisons (ANOVA with Bonferroni correction) for comparison of WT and FGF14 KO inactivation time constants over the range of 0 to ±40 mV were all P<0.01. **(G)** Steady-state inactivation curves (Mean ± SD) following either a 1 sec or 25 ms conditioning step are compared for CCs from WT and FGF14 KO mice. With a 1 sec conditioning step, for WT, V_h_=−50.8±0.7 mV; for FGF14 KO, V_h_=−57.3±0.3 mV. **(H)** Measured voltage of half availability (mean±SD) following 25, 100, 250, and 1000 ms conditioning potentials are plotted for cells from WT (15 cells) and FGF14 KO (14 cells). For comparisons between WT and KO at each conditioning duration, P<0.0001 (ANOVA with Bonferroni correction for multiple comparisons).

We next examined recovery from inactivation first comparing recovery at −80 mV in WT CCs (Fig. 6A) and then in CCs from FGF14 KO mice (Fig. 6B). It can be readily seen that, in the FGF14 KO cell, Nav current is about half recovered within 10 ms, while in WT cells, half recovery only occurs around 100 ms. Similarly, comparison of recovery from inactivation at −120 mV shows that WT cells (Fig 6C) recover much more slowly than FGF14 KO cells. Furthermore, at −120 mV, recovery in FGF14 KO cells is almost complete within 10 ms, while in WT cells recovery is only about half complete in 10 ms. The averaged time course of recovery for CCs from FGF14 KO mice was plotted at voltages from −60 through −120 (Fig. 6E-H) and compared to those from WT mice. The recovery time course for cells from FGF14 KO mice was generally well-described by a single exponential recovery time constant (Fig. 6E-H) in contrast to the double exponentials characteristic of the recovery in WT CCs. However, the average recovery in FGF14 KO cells at −80 mV (Fig. 6F), −100 mV (Fig. 6G), and −120 mV (Fig. 6H) was better fit with the addition of a small, slower recovery component of several hundred milliseconds that contributed a fractional recovery component of about 0.05-0.1. This additional slow component will be addressed below. For the moment, we simply point out that, on average, nearly all of the slow component of recovery from inactivation is abolished when mice do not express FGF14. Furthermore, the remaining fast component of inactivation present in the cells from FGF14 KO mice is indistinguishable from the fast component of recovery observed in WT cells (Fig. 6I). This is consistent with the idea that the fast component simply reflects recovery from inactivation mediated by the normal fast inactivation process that is intrinsic to the Nav1.3 subunit. These results support the view that the entry into the slow recovery pathway predominantly arises from FGF14. Given previous results showing that use-dependent inactivation and slow recovery is produced by FGF14A, but not FGF14B (Laezza et al., 2009; Dover et al., 2010), FGF14A variant is likely the major FGF partner of Nav1.3 in mouse CCs.

**Figure 6.**
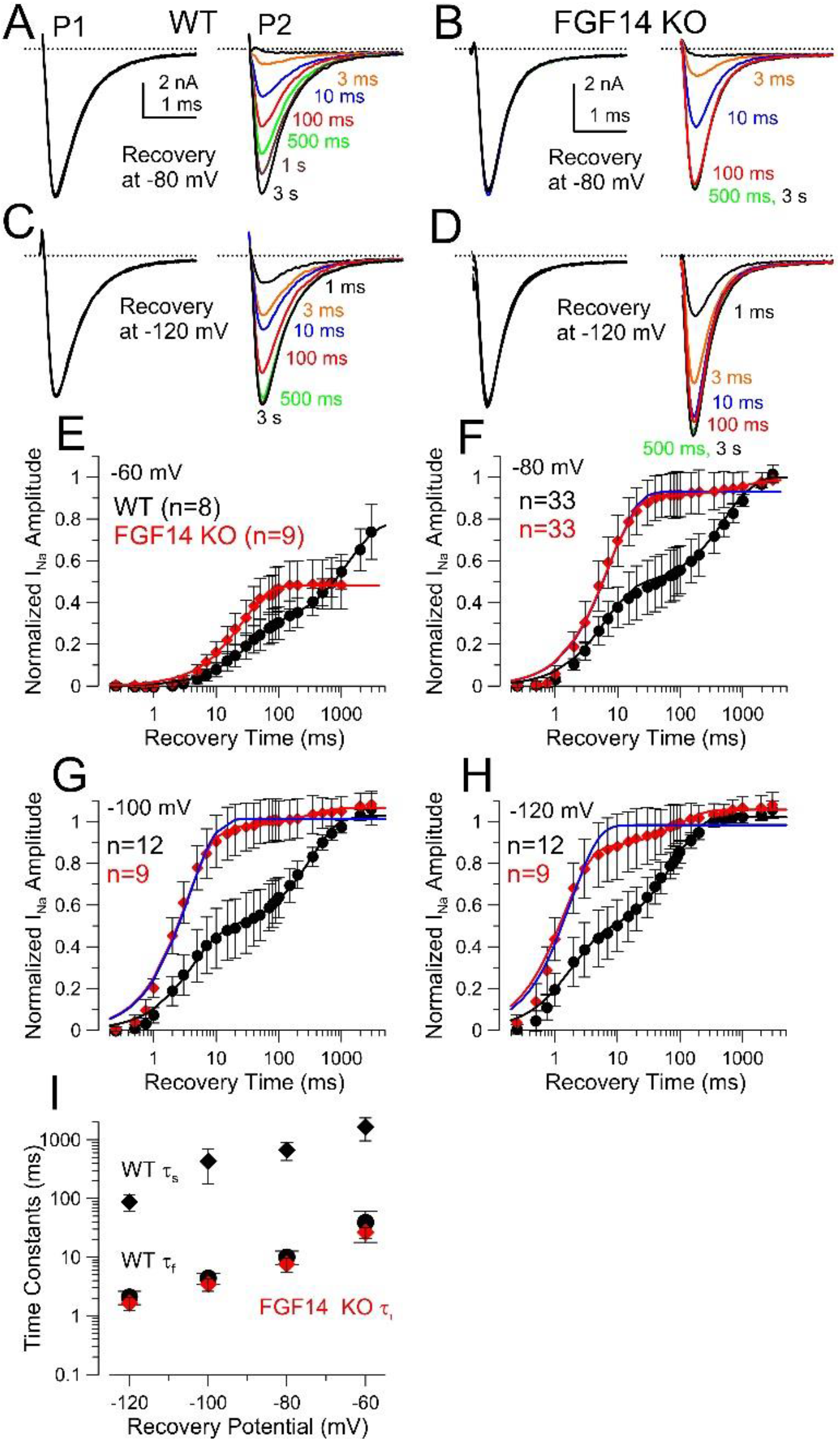
FGF14 KO abolishes all or most slow recovery from inactivation, with little effect on fast recovery. (A) Standard paired pulse recovery at −80 mV for a WT CC. (B) Paired pulse recovery −80 mV for a CC from an FGF14 KO mouse. (C) Paired pulse recovery at −120 mV for the cell shown in A. (D) Paired pulse recovery at −120 mV for the FGF14 KO cell in B. (E) Time course of recovery from inactivation at −60 mV mV (Mean values ±SD) for WT and FGF14 KO CCs (n indicates numbers of cells in all panels). Double exponential fit to WT recoveries was: A_f_ =0.29±0.01, t_f_=32.4±2.7 ms, A_s_=0.49±0.02, t_s_=1336.4±176.0 ms. Single exponential fit to FGF14 KO recoveries: A_f_ = 0.48±0.005 (normalized to full recovery at −80 mV), tf = 26.19±1.07 ms. (F) Recoveries at −80 mV. This includes cells in addition to those in Fig. 2. For WT, A_f_=0.47±0.01, t_f_=6.95±0.59 ms, A_s_=0.53±0.02, t_s_=584±66.3 ms. For FGF14 KO and single exponential fit (blue line), A_f_= 0.92±0.01, t_f_ = 8.40±0.57 ms. For FGF14 KO and two exponential fit (red), A_f_=0.90±0.01, t_f_=7.92±0.46 ms, A_s_=0.10±0.05, t_s_=1129.4±1761.3 ms. (G) At −100 mV, for WT, A_f_=0.48±0.02, t_f_=4.38±0.49 ms, A_s_=0.54±0.02, t_s_=335.3±43.1 ms. For FGF14 KO, single exponential: A_f_=1.02±0.01, t_s_ = 3.83±0.32 ms; for double exponential, A_f_=0.98±0.02, t_f_=3.55±0.30 ms, A_s_=0.09*±*0.04, t_s_=423.4±650.3 ms. (H) At −120 mV, for WT, A_f_=0.46±0.02. t_f_=2.17±0.31 ms, A_s_=0.56±0.02, t_s_=83.3±10.2 ms. For FGF14 KO, single exponential, A_f_= 0.99±0.02, t_f_=2.00±0.28 ms; for double exponential, A_f_=0.89±0.03, t_f_=1.69±0.16 ms, A_s_=0.17±0.03, t_s_=115.9±62.2 ms.(I) Mean (±SD) time constants for slow (ts) and fast (tf) recovery time constants from WT CCs and single exponential recovery time constants for FGF14 KO cells are plotted as a function of recovery voltage.

The similarity of the recovery time constant in the FGF14 KO CCs to the fast component of recovery in the WT CCs indicates that the presence of FGF14 in association with Nav1.3 has little impact on the fast inactivation recovery process. This is consistent with a relative independence of the two inactivation pathways. However, we noted above that the onset of inactivation differed between the WT and FGF14 KO cells (Fig 5E-F). If traditional fast inactivation and FGF14A-mediated fast inactivation are strictly independent, competing inactivation pathways, with rates *k*_f_ and *k*_FGF_, respectively, the expectation is that the inactivation time constant in WT cells would be τ_i_=1/(*k*_f_+*k*_FGF_), while following removal of FGF14A τ_*i*(KO)_=1/*k*_f_. Thus, inactivation following FGF14 KO would be expected to be slower, if onset of inactivation involves two strictly independent, competing pathways. For present purposes, here we assume that all Nav channels contain an FGF subunit. Given that both slow and fast recovery pathways are likely entered at about equivalent rates (equal fast and slow recovery fractions following 5 ms inactivation steps), one would expect the inactivation time constant in the FGF14 KO cells to be about 2-fold slower compared to WT. In contrast, we observed that the onset of inactivation was faster by about 25% (Fig. 5E, at 0 mV) over all tested inactivation voltages (Fig. 5F). Although the magnitude of this difference may not be quite sufficient to approach statistical significance, it deviates from the expectation of a 2-fold prolongation. This result therefore suggests that the traditional fast and FGF14A-mediated inactivation processes may not be entirely independent, but that the presence of FGF14 in the Nav1.3 channel complex may influence the traditional fast inactivation process. The relationship between iFGF-mediated inactivation and traditional fast inactivation will require more detailed evaluation of heterologously expressed Nav currents with and without FGF14A than has yet been done to date.

### Some CCs from FGF14 KO mice exhibit a slow recovery component

As noted above, the averaged recovery from inactivation time course for FGF14 KO cells revealed a small slow component of recovery from inactivation (Fig. 6F-H). Here, we examine the issue of variability in recovery from inactivation among cells of each genotype (WT and FGF14 KO). For WT cells (Fig. 7A), all cells were fit best with a double exponential recovery time course. Although the average amplitude of the fast component was about 0.5 (Fig. 7B), the fast component among individual cells varied from about 0.3 to 0.75. In contrast, for cells from FGF14 KO mice (Fig. 7C), 25 of 33 cells were best fit with a single exponential recovery time constant (cells with black lines in Fig. 7C) while 8 cells (blue) were best fit with a double exponential time course. However, in all FGF14 KO cells for which a double exponential fit was required, the fast component that was in excess of 0.5, markedly different than what was observed in WT cells (Fig. 7A). Furthermore, the slow component of recovery in FGF14 KO cells, when it was present, appeared slower than that in WT cells (Fig. 7G). To better evaluate this behavior, we grouped the FGF14 KO cells into those for which only a single exponential fit was sufficient, and those for which two components of recovery were required (Fig. 7D). Although this separation is somewhat arbitrary, the presence of a slow component in some cells represents a likely physically distinct physical mechanism, rather than a continuum of a single process. For those cells requiring two components (N=8), the fitted values of the averaged recovery were A_f_=0.76±0.01, τ_f_=8.6±0.5 ms, A_s_=0.24±0.06, τ_s_=1450.0±956.6 ms, while for those cells containing only a single exponential component, τ_f_=7.2±0.4 ms. Despite the presence of a distinct slow component in 8 of the set of 33 FGF14 KO CCs we examined and the absence of an explanation for the molecular basis for that slow component, the critical point arising from these observations is that FGF14 is a major determinant of slow recovery from inactivation in essentially all CCs.

**Figure 7.**
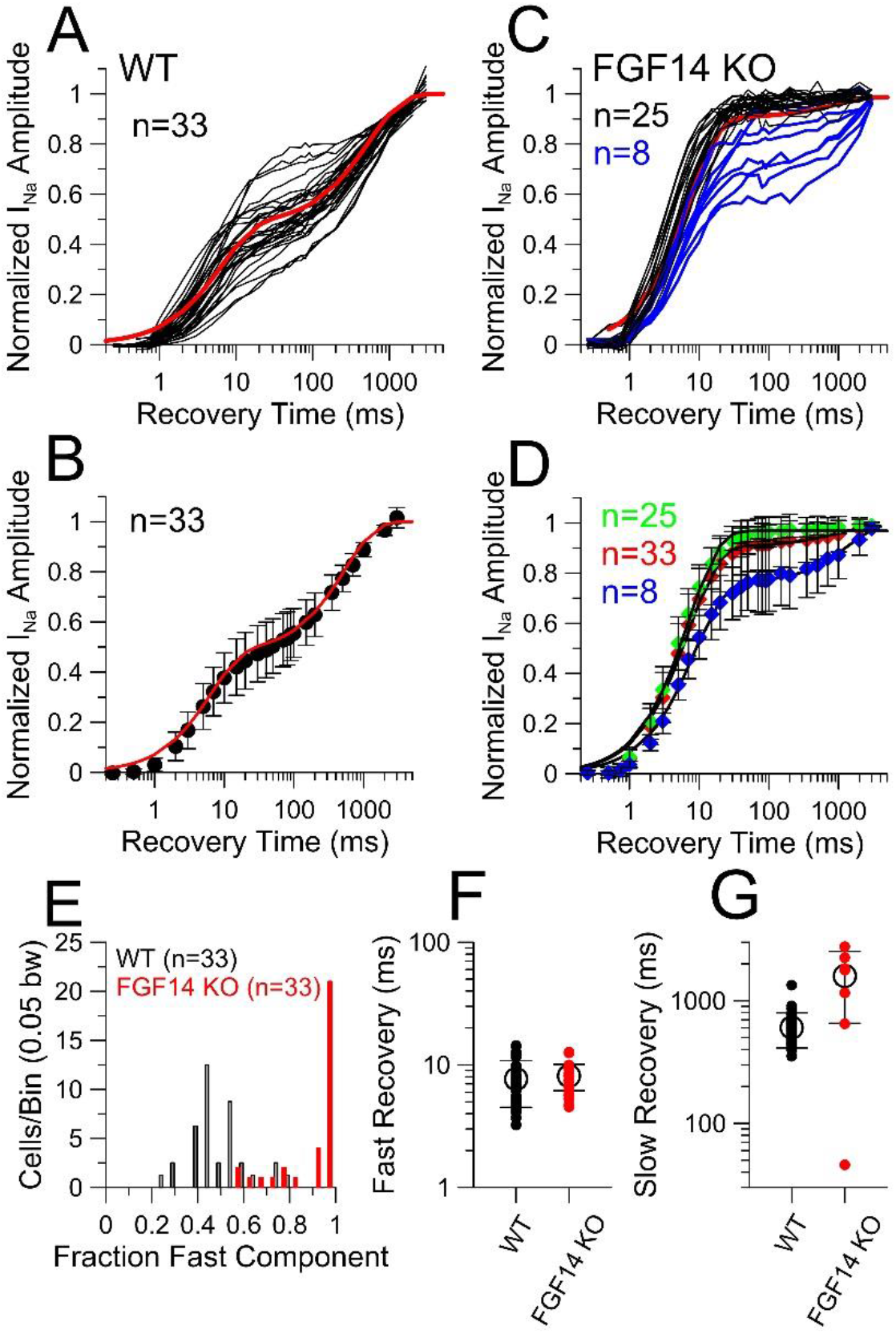
Variability in recovery from inactivation among WT, FGF14 KO, and FGF12 KO cells: some FGF14 KO CCs exhibit a residual component of slow recovery from inactivation. **(A**) Individual fractional recoveries for 16 WT CCs are shown based on the standard paired pulse recovery at −80 mV. Red line shows best fit to averaged recoveries from WT cells. **(B)** The averaged recovery from the 33 cells (from 14 animals) in A, also shown in Fig. 6F. Best fit of a double exponential: A_f_=0.47±0.01, t_f_=7.0±0.6 ms, A_s_=0.53±0.02, t_s_=584.6±66.3 ms. **(C)** Paired pulse recoveries for 33 total CCs from FGF14 KO mice (recovery at −80 mV). Time courses plotted in black were best fit with a single exponential recovery, while those in blue required two exponential components. Red is best fit to averages of all cells. **(D)** For FGF14 KO recoveries, recovery curves are plotted for all cells together (red, identical to FGF14 KO in Fig. 6F), cells fit with only a single exponential time course (green), and cells for which the recovery curve required two exponentials. For grouping of all cells (33 cells from 12 animals) as in Fig. 6F, A_f_=0.90±0.01, t_f_=7.92±0.46 ms, A_s_=0.10±0.05, t_s_=1129.4±1761.3 ms. For FGF14 KO cells in which only a single exponential was required, A_f_=0.97±0.01, t_f_=7.2±0.4 ms. For FGF14 KO cells in which recovery required two exponentials, A_f_=0.76±0.01, t_f_=8.6±0.5 ms, A_s_=0.24±0.06, t_s_=1450.0±956.6 ms. **(E)** Fraction of the fast recovery component for WT (gray) and FGF14 KO (red) were placed in 0.05 wide bins. Fast amplitude values for WT and FGF14 KO were compared using a KS-test because of the non-normal distributions, yielding P=0.000. **(F)** Mean±SD and individual fast time constants are shown for WT and FGF14 KO CCs with adjusted P >0.9999 (ANOVA with Bonferroni multiple comparisons test). **(G)** Mean±SD and individual slow time constants. For WT vs. FGF14 KO, P<0.0001.

The individual values for the fraction of the fast recovery component (A_f_) for WT and FGF14 KO cells were binned and their frequency of occurrence plotted (Fig. 7E). The FGF14 KO (red) cells are clearly shifted towards larger fractions of fast recoveries, even for those cells in which some slow recovery was observed. The fast time constants of recovery from inactivation (Fig. 7F) were indistinguishable between WT and FGF14 KO. In contrast, for those FGF14 KO cells which exhibited slow recovery, the time constants of that slow recovery were slower than those in WT cells (P<0.0001).

### Slow recovery from inactivation is a major determinant of Nav channel availibility

Here we turn to an assessment of the role of slow recovery from inactivation on use-dependent changes in Nav channel availability. In our examination of properties of Nav current in rat CCs, we examined the impact of the slow component of recovery on availability of Nav current during repetitive stimuli (Martinez-Espinosa et al., 2020). The fractional diminution of Nav current either during trains of 5 ms depolarizations or via trains of action potential (AP) waveforms could be readily accounted for by a simple model of two competing, independent fast inactivation pathways. Furthermore, removal of the slow component of recovery from inactivation was predicted to abolish the use-dependent diminution of Nav current amplitude at 10 Hz trains (Martinez-Espinosa et al., 2020). Similarly, for Nav1.5 channels heterologously coexpressed with different FGF13 N-terminal variants (13S,13U, and 13VY), slow recovery from inactivation arising from the FGF13S variant (also termed to FGF13A) is associated with use-dependent diminution of Nav current during 10 Hz trains, while Nav1.5 lacking FGF13 subunits, or those with FGF13U or FGF13VY exhibited no such diminution (Yang et al., 2016). Similarly, diminution of Nav peak current during trains has also been noted for heterologous expression of FGF13A in Nav1.6-expressing ND7/23 cells (Rush et al., 2006).

Here, for set of WT (n=22) and FGF14 KO (n=19) cells (these were part of the set included above (Fig. 7)), we first tested each cell with the standard paired pulse inactivation protocol (1P protocol) with recovery at −80 mV (Fig. 8A, C, E). The same cell was subsequently tested with a protocol (10P protocol) in which a 10 Hz, 10 pulse train of 5 ms depolarizations proceeded recovery intervals at −80 mV (Fig. 8B, D, F). For WT cells, the 10 Hz train drives channels into slow recovery pathways (Martinez-Espinosa et al., 2020), reducing the fast component of recovery from about 0.5 to about 0.3 (Fig. 8G). In contrast, for the FGF14 KO cells we observed two types of behaviors. First, in most cells for which recovery from inactivation was well described by a single exponential (Fig. 8C,D), recovery from inactivation following either the 1P or 10P protocol was essentially identical (Fig. 8H). Furthermore, for such cells, during the 10 Hz train, there was little diminution in the amplitude of Nav current elicited during each pulse in the train (Fig. 8D,J), whereas, in the WT example (Fig. 8B), peak Nav current was reduced to about 40% of the initial amplitude. For those FGF14 KO cells which exhibited a slow component of recovery from inactivation (Fig. 8E-F), the 10 pulse train produced an appreciable decrease in peak Nav current amplitude during the train (Fig. 8F,J), and markedly slowed recovery from inactivation (Fig. 8I). For these three example cells, comparison of the recoveries after the 1P and 10P protocols demonstrate that the WT cell exhibits an appreciable use-dependent increase in the slow component of recovery from inactivation (Fig. 8G), the FGF14 KO cell with no slow recovery shows no difference in recovery between the 1P and 10P protocols (Fig. 8H), while the FGF14 KO cell that exhibited a slow component of recovery shows an even more prominent use-dependent increase in the slow component than in the WT cell (Fig. 8I). It is perhaps noteworthy that in this latter cell, although the slow component of recovery after the 1P protocol is less than the average of WT cells, it exhibits a much more prominent use-dependent increase in the slow component. This is likely based on the fact that, whatever the basis for that slow component of recovery, its time constant is slower than found in WT cells (Fig. 7G). The differences among these cells in terms of the diminution of peak Nav current amplitude during the 10P train are summarized in Figure 8J.

**Figure 8.**
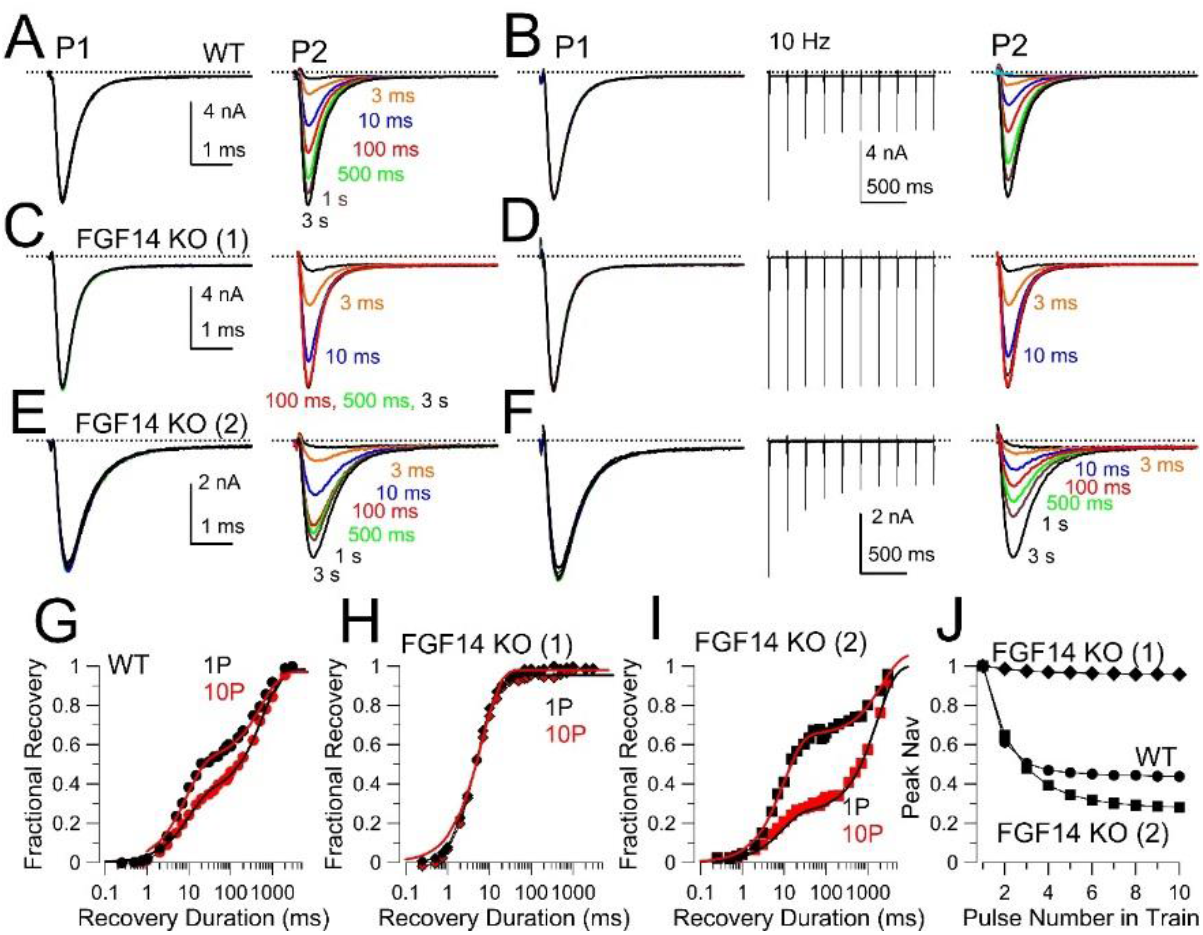
Some CCs from FGF14 KO mice exhibit slow recovery from inactivation. **(A)** A standard paired-pulse protocol was applied to a CC from a WT mouse. On left (P1), overlays of currents elicited by first pulse to 0 mV are shown. On the right, currents evoked by a second test pulse (P2) after recovery intervals at −80 mV are shown. **(B)** For the same cell as in A, recovery was examined after a 10 Hz train of 10 pulses, with P1 and P2 currents as indicated. The variable duration recovery interval was applied at the end of the 10 Hz train. **(C-D)** Identical protocols were applied to CC from a FGF14 KO mouse. Note marked fast recovery (C), and absence of diminution of peak Nav current during train. **(E-F)** Another CC from an FGF14^−/−^ mouse, which exhibited a slow component of recovery from inactivation. Cell was from same tissue slice as that in C-D. **(G)** Fractional recovery for the WT CC following a single step to 0 mV (1P) and following 10 steps to 0 mV (10P). Two exponential fit values were, for 1P, A_f_=0.52±0.01, t_f_=8.9±0.7 ms, A_s_=0.48±0.02, and t_s_=572.6±74.9 ms while, after 10P, A_f_=0.35±0.01, t_f_=9.9±0.8 ms, A_s_=0.63±0.01, and t_s_=604.3±41.9 ms. **(H)** Fractional recovery both following 1P and 10P were fit with single exponentials. Following 1P, t_f_=6.8±0.3 ms, while following 10P τ_f_=7.1±0.5 ms. **(I)** 1P recovery was fit with A_f_=0.65±0.01, t_f_=10.1±0.6 ms, A_s_=0.41±0.12, and t_s_=2252.5±1209.7 ms. After 10P, A_f_=0.28±0.01, t_f_=7.5±0.5 ms, A_s_=0.88±0.06, and t_s_=2379.6±2252.5 ms. **(J)** Diminution in peak Nav current amplitude during the 10P protocol shown in panels B, D, and F for the three cells.

The averaged recovery time courses for the 1P and 10P recoveries for WT (Fig. 9A) and FGF14 KO (Fig. 9B) again highlight the markedly reduced slow component of recovery in the FGF14 KO cells. Individual 10P recoveries for all 22 WT cells (Fig. 9C) and 19 FGF14 KO cells (Fig. 9D) highlight the differences between the WT and FGF14 KO conditions. The 1P and 10P recoveries for each individual cells are plotted for all WT (Fig. S1) and FGF14 (Fig. S2) cells to highlight the generally complete removal of slow recovery in most FGF14 KO cells, but the presence of some slow recovery in three FGF14 KO cells studied with both 1P and 10P protocols. All three of those FGF14 KO cells exhibiting a slow component were from one animal (five additional cells with slow recovery were recorded from five other FGF14 KO animals, but studied only with the 1P protocols; Fig. 7C). It should be noted that in the animal for which multiple cells with slow recovery were observed, other cells from the same animal exhibited a clear single exponential recovery.

**Figure 9.**
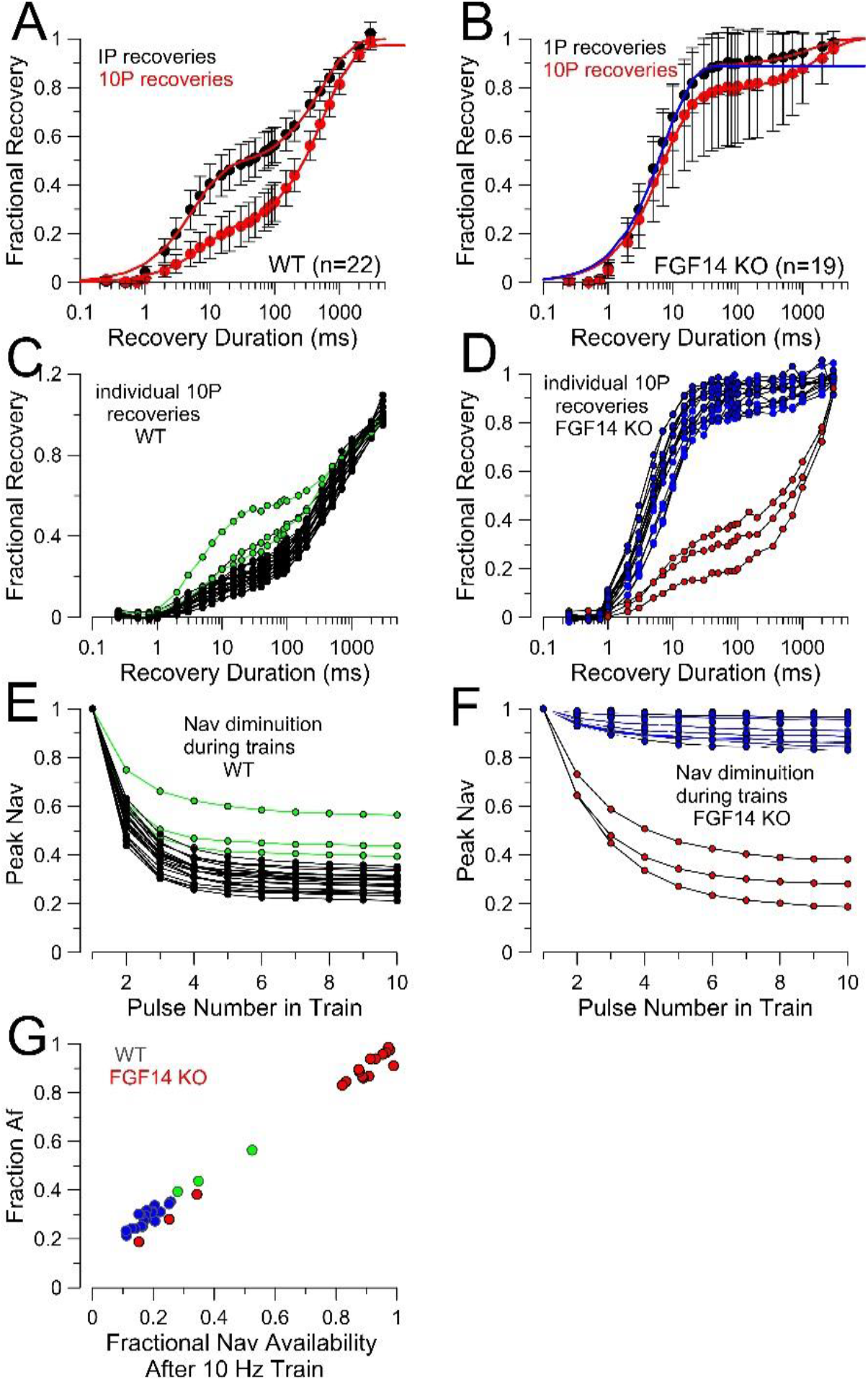
Examination of slow recovery from inactivation in a fraction of FGF14 KO cells. Recovery from inactivation in WT (A) and FGF14 KO (B) cells were compared for recovery at −80 mV from a standard single step (1P) to 0 mV and then for recovery following a 10 Hz train of 10 steps to 0 mV (10P). **(A)** For 22 WT cells, recovery following a 10 Hz train results exhibits a use-dependent decrease in the fast recovery component. For 1P, A_f_=0.47, t_f_=6.03 ms, A_s_=0.53, t_s_=549.4 ms. For 10P, A_f_=0.20, t_f_=7.35 ms, A_s_=0.77, t_s_=589.5 ms. **(B)** For averages of 19 FGF14 KO cells compared with 1P and 10P protocols, the 1P protocol primarily results in exclusively a fast recovery component, but some reduction in the fast component can occur in the 10P protocol (see Fig. S2). Note the large standard error associated with the 10P recoveries. For the 1P protocol, a single exponential fit yielded: A_f_=0.95±0.01, τ_s_=7.5±0.9 ms, while, for a two exponential fit, A_f_=0.94±0.02, t_f_=6.9±0.5 ms, A_s_=0.05±0.07, t_s_=1090.1*±*3892.3 ms. For the 10P protocol, the two exponential fit yielded A_f_=0.79±0.01, t_f_=7.4±0.5 ms, A_s_=0.20±0.1, t_s_=1664.0±1807.2 ms. Note the large confidence limits reflecting the parameters describing the slow component, given that the data do not strongly define this component. **(C)** Recoveries following the 10P protocol for 22 individual WT cells are plotted with the three cells with the largest fast recovery component highlighted in green. **(D)** Recovery following the 10P protocol is plotted for 19 FGF14 KO cells, with the 3 cells showing the most markedly reduced fast recovery highlighted in red. **(E)** Peak current diminution for the 22 WT cells during the 10P protocol are plotted. Green highlights cells in C with a larger component of fast recovery. **(F)** Peak current diminution during the 10 Hz train is plotted for the 19 FGF14 KO cells. For cells with exclusively fast recovery from inactivation (blue), there is little peak current diminution at 10 Hz. **(G)** The fraction of residual fast component (A_f_) following the 10P protocol is plotted as a function of the peak Nav current diminution during the 10P train for WT (blue; green) and FGF14 (red) cells.

Although the limited number of cells and animals for which we have observed the unusual slow component of recovery precludes strong conclusions, a few other features of these cells are worth comment. For the FGF14 KO cells that exhibited a slow recovery component following 1P stimulation, the slow fraction is smaller than that typically seen in WT cells. However, recovery following the 10P protocol is associated with a slow component of comparable amplitude to that seen in WT cells (Fig. 9D), but of slower duration (Fig. 8I). We would suggest that this strong use-dependent increase in the slow recovery fraction despite a modest component of slow recovery following a single inactivation step results in a somewhat more slowly developing, but similarly profound use-dependent diminuition in peak Nav current during the 10 Hz train of stimuli, compared to WT cells (Fig. 9E vs. 9F). Our results provide no direct test of the basis for the slow component of recovery in some FGF14 KO cells, but we anticipate that generation of KO mice for other FGF isoforms may be informative on this issue. To highlight the relative importance of the accumulation of channels in slow recovery pathways on use-dependent changes in peak Nav amplitude, we plotted the fraction of the fast recovery component following the 10 Hz train versus the fractional Nav availability based on measurement of relative Nav current amplitude during the 10^th^ pulse in the train (Fig. 9G). Although most FGF14 KO cells exhibited a large fast amplitude component along with little diminution in Nav availability, for the FGF14 KO cells with a slow component of recovery, both the reduction in fast amplitude and reduction in Nav availability mirror the behavior seen in WT cells. Furthermore, three WT cells exhibiting the largest fast recovery component after the 10P protocol (green in Fig. 9C,9E,9G) exhibited the smallest use-dependent diminution of peak Nav current. Thus, it is the accumulation of channels in slow recovery pathways that dictates the extent of diminuition of peak Nav current during trains.

### Dual-pathway inactivation and use-dependent changes in Nav availability

In the associated paper, a simple set of assumptions about the dual-pathway inactivation process described well the use-dependent diminution in Nav currents activated either by trains of square pulse depolarizations or action potential voltage-clamp waveforms (Martinez-Espinosa et al., 2020). The basic assumptions were the following: 1. during a 5 ms depolarization or an AP, all activated channels inactivate with half entering traditional fast recovery states and half entering a slower recovery pathway; 2. following inactivation, channels in either recovery pathway recover from inactivation into resting states in accordance with experimentally measured time constants. Therefore, beginning with 95% of more of the channels in resting states (assuming a −80 mV holding potential), fractional occupancy prior to any test pulse, immediately following any test pulse, and then after the recovery interval preceding the next test pulse can be readily calculated (Fig. 10A). Using this approach and the measured fast and slow time constants of recovery from inactivation at −80 mV, the occupancy of channels in closed, fast inactivated, or slow inactivated states was determined for the WT condition at a time preceding each test depolarization in a 20 pulse train at 10 Hz (Fig. 10B1), and also following each 5 ms test depolarization where all channels are assumed to be inactivated (Fig. 10B2). From calculation of the channels that have returned to closed states preceding each depolarization, the calculated fractional decrease in peak Nav current amplitude was determined for the given set of recovery constants (Fig. 10B3). Finally, based on the occupancies of channels in fast and slow recovery pathways following either a single test step (1P) or a train of 10 pulses (10P), the predicted 1P and 10P fractional recovery from inactivation was determined (Fig. 10B4). Both the diminution of Nav amplitude and the relative changes between 1P and 10P recovery time courses match well with the observations above. To illustrate the impact of FGF14 KO, in which no slow component of recovery from inactivation was observed, we repeated the calculations for a case in which only fast inactivation is observed (Fig. 10C1-C4). As trivially expected, there is no diminution of peak Nav amplitude during the train (Fig. 10C3) as also observed experimentally (Fig. 8D). Finally, we considered a case analogous to that of FGF14 KO cells in which a smaller fraction of initial slow recovery is also associated with a slower recovery time constant, using the experimentally measured values of A_f_=0.75, and τ_s_=1591 ms (Fig. 10D1-D4). That 75% of channels inactivate into fast recovery pathways, an empirically based assumption, could arise from a number of mechanistically distinct factors, about which we have no information. However, employing those values, peak Nav current amplitude can be seen to diminish to values comparable to the WT case, although with a slower approach to steady-state (Fig. 10D3). Similarly, the calculated 1P and 10P recoveries show the larger fractional increase in the slow recovery component (Fig. 10D4) observed in those FGF14 KO cells, which exhibited a slow recovery component. These considerations suggest that differences in the time constant of slow recovery in dual pathway system can profoundly impact on the rate of use-dependent accumulation in slow recovery and, correspondingly, temporal aspects of changes in Nav availability during trains of stimuli.

**Figure 10.**
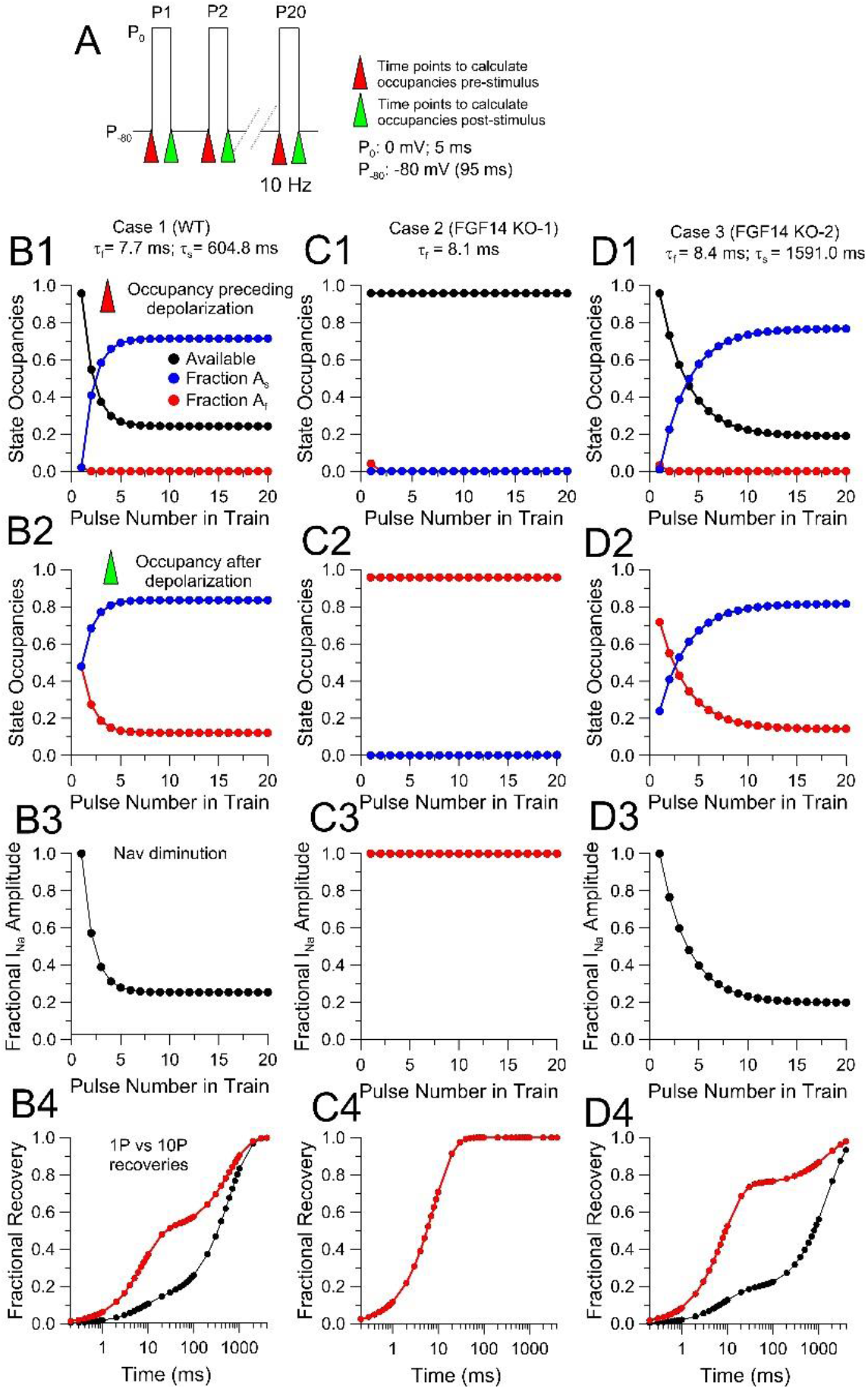
Evaluation of the impact of slow and fast recovery from inactivation on use-dependent diminution of peak Nav amplitude. **(A)** The basic stimulation protocol for calculation of state occupancies, use-dependent peak current diminution and 1P/10P recovery time course. From a −80 mV holding potential, a 10 Hz train of twenty 5 ms steps to 0 mV was used to produce channel activation and inactivation. This corresponds to a 95 ms recovery interval between steps to 0 mV. Red triangles indicate time points prior to depolarization and green, after full inactivation. For calculations in B-D, fractional inactivation prior to P1 is assumed to be 0.05, with inactivated states split evenly between fast and slow recovery pathways. **(B)** Calculations assume channels available for activation at beginning of depolarization equally enter slow and fast recovery pathways, with no interconversion. Fraction of slow and fast recovery during each 95 ms recovery interval is calculated based on measured time constants of fast and slow recovery for WT cells at −80 mV (Fig. 7F-G) and calculated initial fraction of channels in either fast or slow pathways. **B1**, black circles indicate fraction of channels available for activation prior to depolarization, red indicates channels in fast recovery pathways prior to depolarization, and blue indicates channels in slow recovery pathways prior to depolarization. **B2**, fractions of channels in slow recovery pathways (blue) and fast recovery pathways (red) immediately following inactivation (Green arrows in A). **B3**, calculated fraction of peak Nav current normalized to P1 amplitude based on channels available for activation prior to each test pulse to 0 mV. **B4,** time course of recovery from inactivation based on calculated fraction of channels in fast and slow pathways following P1 or P10 and the measured time constants of recovery. **(C)** Calculations similar to those in B, but assuming that the slow component of recovery from inactivation is absent (most FGF14 KO cells. This yields no appreciable use-dependent diminution in Nav amplitude and identical single exponential time courses of recovery from inactivation following P1 or P10. **(D)** Calculations as in B and C, but assuming average fast and slow recovery time constants based on the set of FGF14 KO cells that exhibited a slow component of recovery.

To probe the potential roles of different aspects of the dual pathway model on use-dependent changes in Nav availability, we first tested the impact of varying the fractions of channels entering A_f_ and A_s_ with each inactivating step (Fig. 11A). This might arise because intrinsic rates of traditional fast inactivation and FGF-mediated inactivation differ. For example, the observed macroscopic inactivation time constant of about 0.4 ms at 0 mV (Fig. 1E) would be consistent with a range of combinations of intrinsic rates of entry into fast and slow pathways, all summing to 2500/s, where τ_i_=1/(*k*_i_+*k*_FGF_), where A_f_=*k*_i_/(*k*_i_+*k*_FGF_), where *k*_i_ is intrinsic rate of traditional fast inactivation and *k*_FGF_ is rate of inactivation via FGF-mediated inactivation. A second factor affecting A_f_ and A_s_ could be less than full stoichiometric presence of an FGF subunit in the Nav population. However, for the present calculations, this latter possibility is not considered. We utilized rates of fast and slow recovery from inactivation identical to those measured in WT CCs. The rates of recovery from inactivation following a single pulse (Fig. 11A1) simply mirror the fraction of occupancy dictated by the assumption of a given A_f_ value. The diminution of peak Nav current varies considerably as the relative entry into fast and slow recovery pathways varies (Fig. 11A2) with the most dramatic effects on cumulative inactivation seen over A_f_ values from 0.5 to 0.95. Furthermore, the rate at which a steady-state level of Nav availability is attained varies considerably. As A_f_ increases, this naturally slows entry into slow recovery pathways during a train of stimuli. The accumulation of channels in slow recovery pathways is highlighted with calculation of recovery following a standard 10P protocol (Fig. 11A3). The impact of varying the time constant of recovery from fast inactivation from 6 to 30 ms was also evaluated (Fig. 11B). In this case, although the shape of the recovery from inactivation following a 1P protocol (Fig. 11-B1) varies as expected for changes in τ_f_, τ_f_ has no impact on the use-dependent accumulation of Nav channels in slow recovery pathways, as revealed in either the Nav peak current (Fig. 11-B2) or the 10P recovery protocol (Fig. 11-B3). When the slow time constant of recovery (τ_s_) is varied from 200 through 3000 ms (Fig. 11-C1), slowing of ts results in a marked increase in the extent of use-dependent diminution of peak Nav current during a train (Fig. 11-C2), but with only modest impact on how much diminution occurs with each stimulus during a train. The latter effect, in contrast to what was observed with different values of entry into slow and fast recovery paths (Fig. 11-A), arises since τ_s_ does not impact on how many channels enter slow recovery states with each stimulus, but only the persistence of channels in the slow recovery rates between stimuli. The 10P protocol also emphasizes the impact of τ_s_ on the shape of the time coruse of recovery from inactivation (Fig. 11-C3). Finally, we again examined the consequences of varying A_f_ (differential entry into fast and slow recovery pathways), but in association with a recovery time constant similar to what we observed in those FGF14 KO CCs with a slow component of inactivation (Fig. 11-D). Although the 1P recovery behaviors are generally similar to those with a faster τ_s_ (Fig. 11-D1), the calculations of peak Nav current diminuition during a train further highlight the impact that a given τ_s_ can have on the approach to steady-state availability during a train particularly in conjunction with reduced fraction of entry (e.g., A_f_ = 0.95) into slow recovery pathways (Fig. 11-D2). The greater accumulation of channels in slow recovery pathways is also highlight in the 10P protocol (Fig. S11-D3).

**Figure 11.**
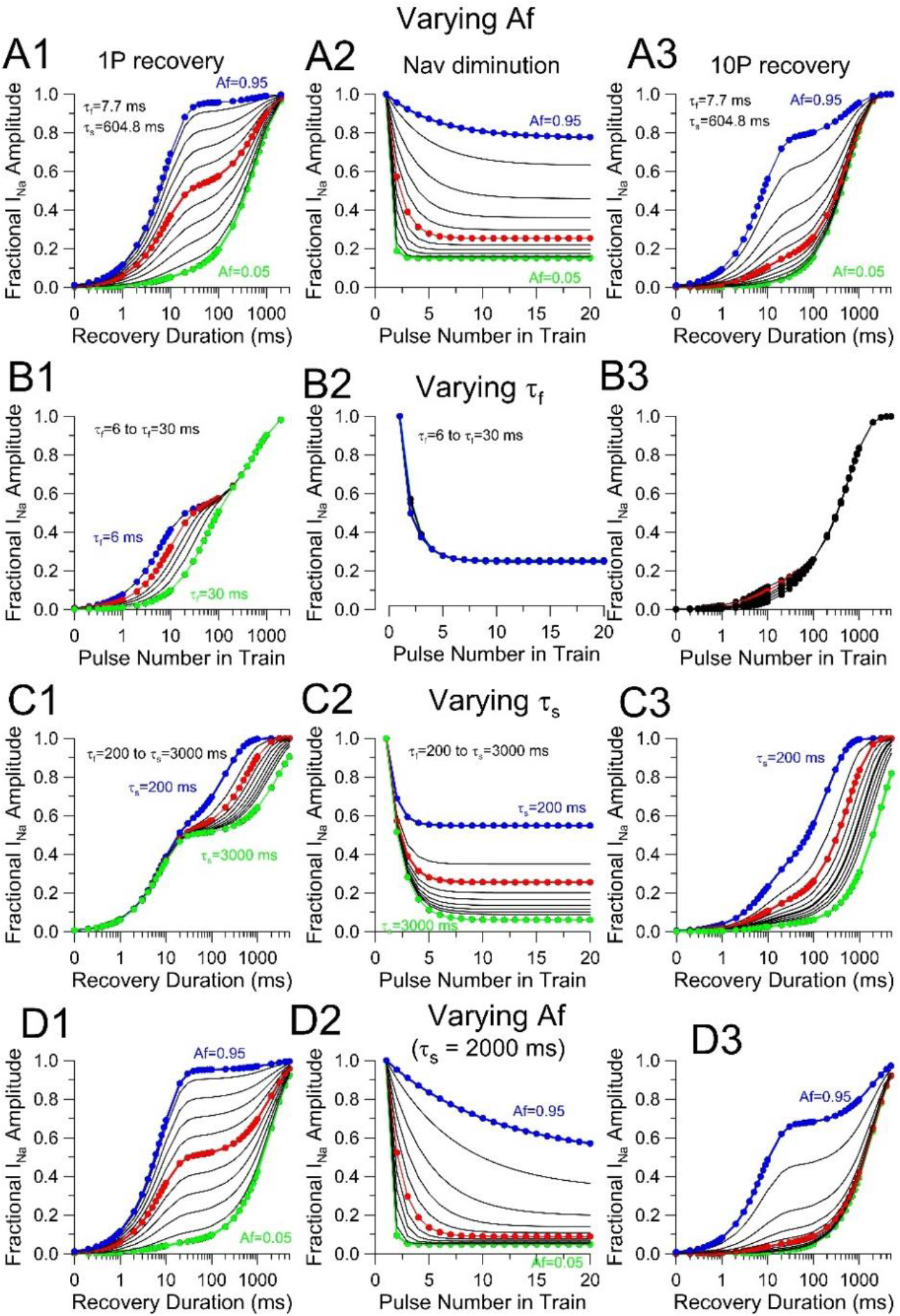
Impact of changes in inactivation properties on predicted Nav diminution during trains and increases in slow recovery fraction. Underlying assumptions are summarized in the text in association with Fig. 10. Based on Fig. 5F, fast inactivation in absence of FGF is about 0.4 ms. Even if entry into slow recovery is 10-fold slower than normal fast inactivation, it is expected that inactivation will be complete during a 5 ms inactivation step, but that fewer channels will be in slow recovery pathways. (A) Impact of varying A_f_ (A_s_=1-A_f_) on predicted 1P recovery time course (A1), Nav diminution extent and time course (A2), and 10P recovery (A3). Blue: A_f_=0.95; red: A_f_ = 0.5; green: A_f_=0.05. Red corresponds to WT CC behavior. (B) As in A, but with predictions based on varying tf from 6 ms (blue) to 30 ms (green), with other constants as for WT. (C) As above, but with predictions based on varying ts from 200 ms (blue) to 3000 ms (green) with red similar to to WT measurements (600 ms). (D) Predictions based on variation in A_f_ from 0.95 (blue) to 0.05 (green), but with ts = 2000 ms, highlighting profound impact of τ_s_ on cumulative inactivation (D2 and D3)

The main point to be made from the above considerations is that, if there are specific molecular considerations that can influence either 1. relative rates of entry into slow and fast recovery pathway or 2. rates of recovery via the slow recovery pathway, such factors would dramatically impact on use-dependent changes in Nav current availability during repetitive activity. Although a major portion of the use-dependent changes in Nav availability in mouse CCs clearly arises from the presence of FGF14 subunits, the presence of a slow recovery behavior of different properties (e.g., reduced slow component following a single depolarization and different τ_s_) at least in some FGF14 KO CCs suggests there may be other factors, perhaps different FGF subunits, which can sculpt changes in Nav availability among different cells.

## Discussion

The present results establish that the Nav1.3 isoform of sodium channel accounts for all Nav current in mouse chromaffin cells studied in adrenal medullary slices. Furthermore, FGF14 appears to be an integral component of all (or most) mouse CC Nav channels, being the primary determinant for the slow component of recovery from fast inactivation in most cells. This may the first case for which an FGF is shown to partner with Nav1.3 in a native tissue, although, given conservation of the FGF-binding surfaces among Navs, that is not surprising. Here, we will discuss four main issues: first, the relevance of the present observations to earlier work on Nav currents in CCs, second, possible explanations for slow recovery from inactivation that is present in a subset of CCs from FGF14 KO mice, third, implications of the present work concerning independence of traditional fast inactivation in relationship to FGF-mediated inactivation, and, finally, the potential impact of FGF-mediated inactivation and slow recovery from inactivation on Nav availability and excitability.

### Molecular substrates of Nav currents in CCs

Given the similarity in properties between rat and mouse CC Nav currents, particularly in the steady-state inactivation behavior, it must be considered that the Nav currents in rat CCs also arise from Nav1.3. However, in the absence of tests that might fully knock-down Nav1.3 expression in rat CCs, this point remains uncertain. That Nav1.3 is the predominant Nav current in mouse CCs runs counter to the long-standing view that Nav1.7 is the dominant neuroendocrine sodium channel (Wada et al., 2004; Yanagita et al., 2007; Wada et al., 2008). Much of this earlier body of work focused on bovine CCs and examined the up- or down-regulation of Nav1.7 without assessing the potential contribution of other Nav isoforms.

Evidence supporting a role for Nav1.7 is summarized here. Both Nav1.7 message and presumed protein has been found to be present in dissociated bovine CCs (Wada et al., 2004; Yanagita et al., 2007; Wada et al., 2008). Protein was defined by saxitoxin binding, which is not definitive for Nav1.7, and also the use of monoclonal antibody NS68/6, Nav1.7 protein, without KO controls. Whether message for other Nav isoforms may be present in bovine CCs has not been described, to our knowledge. To date the only recordings of Nav currents in bovine CCs appear to be those from an early seminal paper (Fenwick et al., 1982) and over the years it appears to have simply been assumed that such currents likely arise from Nav1.7. Is there anything to suggest this is not the case? The steady-state inactivation curve of bovine Nav currents using a 25 ms conditioning step resulted in a V_h_ of about −35 mV (Fenwick et al., 1982), which is comparable to the rat steady-state GV measured after a similar 25 ms conditioning step. When the GV in bovine cells was measured from two holding potentials, −80 and −100 mV, currents following the −100 mV holding potential resulted in an approximately 10% increase in peak current activation. This is quite different for one unambiguous case where both Nav1.3 and Nav1.7 have been shown to be expressed in the same cells (Zhang et al., 2014). Specifically, large differences in holding potential (−70 vs. −180 mV) prior to steady-state inactivation protocols have been productively used with mouse pancreatic β and α cells to reveal two distinct components of Nav current, Nav1.3 and Nav1.7, highlighting that at −180 mV a component of current corresponding to Nav1.7 is available for activation, but which is almost completely inactivated at −70 mV (Zhang et al., 2014). Although additional tests would be required in CCs, we would suggest that the additional Nav current unmasked in bovine CCs with a holding potential of −100 mV (compared to −80 mV) may reflect Nav1.7 in such cells, with perhaps Nav1.3 making the more substantial contribution to bovine CC Nav current. For the pancreatic β and α cells, the use of Nav1.3 and Nav1.7 KO mice confirmed the identities of the more left-shifted current component (Nav1.7) and right-shifted component (Nav1.3) (Zhang et al., 2014). A similar approach comparing holding potentials of −120 mV or −70 mV on steady-state current availability in rat pancreatic β cells and rat CCs revealed that, whereas there is little difference in CC Nav current between the two holding potentials, in rat β cells almost all Nav current is abolished from a holding potential of −70 mV (Lou et al., 2003). Given the idea that the steady-state availability of Nav1.7 is substantially shifted leftward compared to Nav1.3, these latter results also support the idea that Nav1.7 is predominant in rodent pancreatic β cells, but that Nav current in rat CCs is likely to arise from Nav1.3. Although steady-state inactivation can be influenced by auxiliary subunits, at least for Nav1.3 none of the known β subunits appear to shift steady-state inactivation close to properties of Nav1.7 channels, when they are expressed heterologously (Cummins et al., 2001).

Overall, the present results support the idea that the functional properties of Nav current in both mouse and perhaps rat CCs arise from a single variety of Nav channel. This appears to be true despite the presence of both Nav1.3 and Nav1.7 message in rat (seen here) and, to a lessor extent, in mouse where Nav1.7 message is less abundant (Vandael et al., 2015) and the present results). Clearly, the availability of low-affinity (reversible), selective blockers of different Nav isoforms would help address this issue. Given the generally left-shifted gating behavior of Nav1.7 currents relative to Nav1.3 (Zhang et al., 2014) and the usually complete inactivation of Nav1.7 at potentials positive to about −60 mV, it is unclear what physiological role any Nav1.7 current in chromaffin cells might play, if they were present.

### FGF14A is likely the major determinant of the slow recovery from inactivation in mouse CCs, but other contributors may exist

The slow recovery from inactivation observed in mouse CCs arises primarily from the FGF14 subunit, which we suggest likely reflects specifically FGF14A. The complete absence of slow recovery from inactivation in most CCs from the FGF14 KO mice argues that FGF14A is an essential partner in the Nav1.3 channel complex in CCs and is the primary determinant of the slow component of recovery from fast inactivation. However, we also found a subset of FGF14 KO cells for which a slow recovery component was observed. This additional slow component was, on average, smaller in relative amplitude and slower in time course than in WT cells, indicative that the molecular basis of the persistent slow recovery is distinct from that arising from FGF14.

What are potential explanations for the presence of the residual slow recovery in some FGF14 KO cells? For the 8 of 33 FGF14 KO cells exhibiting some slow recovery from inactivation, on average the initial fraction of slow recovery following a single inactivation step was 0.25 with an average time constant of slow recovery about 2-fold slower than in WT cells. One explanation, which we consider the most likely, is that this component may arise from the presence of another, as yet unidentified, FGF isoform. This isoform, although perhaps in lower abundance, may contribute along with FGF14 in WT cells in influencing Nav inactivation behavior. However, in WT cells, it would be virtually impossible to separate out multiple distinct slow recovery processes. This possibility will require evaluation with KO animals for additional FGF isoforms, when they are available. This possibility has several intriguing implications which must also be evaluated in future work. First, the slower time constant of recovery of this persistent component suggests that this additional FGF subunit has kinetically distinct effects from FGF14A on Nav1.3 channels. Not only is the rate of recovery from FGF-mediated inactivation slower, but the smaller fraction of initial slow pathway occupancy following a single inactivation step suggests that the onset of inactivation mediated by the additional component may be slower. Second, the kinetically distinct behavior of the slow recovery occurring in some FGF14 KO cells was shown to be associated with a slower, but equally deep development of use-dependent diminution of peak Nav current. This suggests that, if the slow component observed in some FGF14 KO CCs does arise from an additional FGF isoform, use-dependent changes in Nav variability may be differentially regulated by expression of particular FGF subunits. Thus, functional differences among FGF-A isoforms have the potential to differentially impact on use-dependent changes in Nav availability. An alternative explanation of the slow recovery of some FGF14 KO cells is that it arises from something other than an FGF subunit. However, given that the behavior of the slow component mirrors aspects of FGF14, we would not favor this possibility.

A few other points regarding the FGF14 cells in which slow recovery was observed must be mentioned. Of the 8 FGF14 KO cells for which slow recovery were observed, three were from the same animal. However, an important point is that other cells from the same animal exhibited exclusively fast recovery from inactivation. Thus, the residual slow recovery in FGF14 KO animals occurs in only some CCs within a given animal, suggesting heterogeneity in expression of the determinant of the additional slow recovery process. Although we have only a limited number of such cells, slow recovery from inactivation was also observed in CCs from both male and female FGF14 KO mice.

### The relationship between FGF-mediated inactivation and traditional fast inactivation

Since the initial work of Goldfarb and associates, the prevailing hypothesis regarding dual-pathway fast inactivation (or long-term inactivation) is that traditional fast inactivation intrinsic to the Nav channel subunit and inactivation mediated by the A-isoform of an FGF compete to occupy overlapping positions of occupancy that occlude the ion permeation pathway (Dover et al., 2010; Goldfarb, 2012; Venkatesan et al., 2014). Based on current thinking, fast inactivation involves a peptide loop containing a conserved IFM fast inactivation motif on the linker between domains III and IV of inactivating Nav channels. Although at one time the IFM loop was thought to act as a cap to hinder ion flux through the permeation pathway (West et al., 1992; Eaholtz et al., 1994), current structural evidence suggests that it may act allosterically to constrict ion flux (Yan et al., 2017). Furthermore, it is movement of the Domain IV voltage sensor that is sufficient for fast inactivation in sodium channels and also limits the rate of fast inactivation (Capes et al., 2013). For intracellular FGFs, a conserved homologous core domain present in all FGF isoforms has been shown to mediate binding to Nav alpha subunits and mutations of that conserved surface disrupt the ability of FGFs to assemble with Navs (Goetz et al., 2009; Wang et al., 2012). The present understanding of how FGF A-isoform N-termini produces inactivation has been largely based on functional tests of coexpression of FGF13A with Nav1.6 (Dover et al., 2010). This work (Dover et al., 2010) demonstrated that the N-terminus of the FGF13A protein produces long-term inactivation of Nav1.6 channels, that mutation of basic residues in the N-terminus abolishes the long-term inactivation, and synthetic peptides corresponding to residues 2-21 of the FGF13 subunit produces use-dependent inhibition of Nav1.6 current in a fashion similar to inact FGF13 subunits. In contrast, coexpression of FGF13B with Nav1.6 does not produce long-term inactivation. Together, these results led to the proposal that the mobile cytosolic N-terminus of the FGF A-isoforms could occlude ion permeation, in a fashion that competes with the normal fast inactivation process, but involves a slower recovery from inactivation (Dover et al., 2010; Goldfarb, 2012). Although such evidence is consistent with channel occlusion models developed for fast inactivation of K^+^ channels, as noted for the case of the IFM motif mediating fast Nav channel inactivation, the specific molecular unpinnings of the FGF-mediated inhibition of ion permeation remain to be determined.

The present results and the associated paper (Martinez-Espinosa et al., 2020) address a few points pertinent to the relationship between the two inactivation pathways. First, results are generally consistent with the idea of independence between the two pathways. Namely, once inactivation into one or the other pathway has occurred at strongly inactivating conditions, channels appear effectively precluded from equilibrating into the other pathway. The use-dependent accumulation in slow recovery pathways also strongly supports the idea that the two pathways are largely independent. However, one aspect of our results conflicts with this idea. Specifically, the onset of inactivation in CCs from FGF14 KO mice is more rapid than in CCs from WT mice (Fig. 5E-F) and a simple shift in the activation GV is unlikely to account for these differences. For FGF14A-mediated fast inactivation in mouse CCs, our results indicate that about half the channels inactivate into each pathway. Assuming all channels contain an inactivating FGF subunit, this requires that the rates of entry into each are quite similar. If two independent fast inactivation mechanisms with comparable rates defined by *k*_f_ and *k*_FGF_ compete to produce inactivation, the expected time constant of inactivation would be defined by τ_i_=1/(*k*_f_+*k*_FGF_). Since at strong inactivation voltages entry into either inactivated state is largely absorbing, only a single exponential is expected. If the two pathways are independent, in the absence of FGF14, inactivation should then be defined simply by τ_i_=1/*k*_f_ which predicts a time constant twice as slow as in the WT. Thus, that inactivation is actually faster in CCs from the FGF14 KO mice raises the possibility that, at least for entry into inactivated states, there may be some interaction between the two pathways. This might reflect an allosteric effect or simply a steric effect whereby, for example, the presence of FGF14A, might hinder but not prevent movement of the IFM-containing DIII-DIV linker. Such questions can most effectively be addressed with heterologous expression studies. But this potential nuance of FGF14A/Nav interaction does not contradict the key idea that, in large measure, the two pathways are largely independent.

### FGFs, steady-state inactivation, slow recovery from inactivation, and cell firing

There is an extensive literature concerned with various aspects of the specificity of interactions of different FGF isoforms with different Nav isoforms (Liu et al., 2003; Goetz et al., 2009; Laezza et al., 2009; Wang et al., 2012) and potential Nav/FGF partnerships in native tissues (Bosch et al., 2015; Wang et al., 2017; Effraim et al., 2019). Furthermore, there has been extensive attention to the physiological and pathophysiological consequences of different FGF isoforms and naturally occurring mutations (Liu et al., 2003; Laezza et al., 2007; Hennessey et al., 2013; Musa et al., 2015; Siekierska et al., 2016). Many of the functional studies have largely focused on the impact of FGFs on activation GVs, steady-state inactivation curves, channel localization, or current density (Goldfarb et al., 2007; Diwakar et al., 2009; Xiao et al., 2013; Pablo et al., 2016; Yang et al., 2016), suggesting that shifts in steady-state inactivation curves, by changing Nav channel availability at resting potentials, may impact on cell firing. When examined in heterologous expression systems, both A and B FGF isoforms appear to produce generally similar shifts in steady-state inactivation. There is no question that shifts in steady-state inactivation among cells will differentially impact on Nav availability, but such shifts *per se* are unlikely to underlie use-dependent alterations in availability, unless associated with specific kinetic behaviors that result in accumulation of channels in states that only slowly recover from inactivation.

Thus, it is the use-dependent accumulation in slowly recoverying inactivated states which is the main determinant of use-dependent diminution of Nav channel availability during trains, in line with initial proposals from Goldfarb (Dover et al., 2010; Goldfarb, 2012; Venkatesan et al., 2014). Invariably, where it has been examined with heterologous expression, such use-dependent diminution of Nav channel availability has been specifically associated with FGF-A isoforms, for which slow recovery from inactivation is also observed. This has been shown for FGF14A coexpression with Nav1.6 channels (Laezza et al., 2009), for FGF-mediated spike accommodation in hippocampal pyramidal cells (Venkatesan, 2014 #9396}, for FGF13S-mediated coexpression with Nav1.5 channels (Yang et al., 2016), and for FGF13A coexpression with Nav1.6 (Rush et al., 2006). The extent of the steady-state diminution of peak Nav current shows considerably variability among the various cases ranging from about 50% reduction to about 90%, even at identical 10 Hz train frequencies. Ultimately, as supported by analysis here the specific kinetics of the FGF-mediated onset and recovery process are likely to be essential in defining the extent and time course of use-dependent diminution of Nav currents, but at present the conditions under which slow recovery has been examined in work to date prevent any generalities to be drawn regarding different FGFs and different Nav channels

An important pair of papers on Nav currents in dorsal raphe neurons (Milescu et al., 2010; Navarro et al., 2020), almost certainly involving some as yet unidentified FGF, provides some of the more useful comparative information to the results presented here. Interesting differences between the Nav currents in CCs and those in dorsal raphe are that, whereas in CCs, inactivated channels are about equally distributed between fast and slow recovery pathways during a single depolarizing step, in dorsal raphe neurons only about 20% of channels enter initially slow recovery pathways (Milescu et al., 2010; Navarro et al., 2020). At present, we lack sufficient quantitative information about the differential entry and rates of recovery among different cell types to draw any conclusions, but below we consider a few factors that will need to be considered in future work.

#### Are most Nav channels in native cells partnered with an FGF?

Based on the known binding interaction surfaces for Nav isoforms and FGF subunits, the functional effects of FGF’s probably involve the presence of only a single FGF molecule in the complex. Yet, in native cells we can not as yet be certain how well-populated the Nav C-terminal FGF binding motif may be. The measurement we have available is the initial fractions of channels in fast and slow recovery pathways following brief inactivation steps, which for both rat and mouse CCs is about 0.5 for each. As mentioned, this does not appear to be case for raphe neurons (Navarro et al., 2020). For most other experimental work on FGFs to date, such quantitative information is not available. Although we have no direct information regarding the rates of entry into each recovery pathway in CCs, the equal distribution between slow and fast pathways would suggest a similarity in such rates at least at 0 mV and above. However, this assumes that all Nav channels in a cell are associated with FGF subunits about which we have no direct information. Can we draw any inferences regarding the extent to which all Nav channels in CCs may partner with an FGF? Experimentally, the fact that perhaps 80-90% of channels can be driven into slow recovering pathways, at least in some cases, gives some confidence that most Nav channels are partnered with an FGF.

However, there are limitations in the use of the initial fractions of fast and slow recovery populations as an indicator of extent of FGF assembly in a channel population. First, for those FGF14 KO cells which exhibited a slow component of recovery, the initial fraction of slow recovering channels was typically 0.25 or less, being about 0.4 at most (Fig. S2), suggesting perhaps less assembly of any putative additional FGF in the channel population. Yet, in those cases the 10P protocol could still drive as much as 0.8 of the channel population into slow recovery pathways, indicative that, despite a modest initial fraction of entry into slow recovery pathways, most (or perhaps all) Nav channels still might contain a subunit mediating such slow recovery. This might arise, if rates of initial entry into an FGF-mediated inactivated state were substantially slower than into traditional fast inactivated states at 0 mV for a given type of FGF subunit. Second, in our calculations addressing the impact of different fractions of entry into fast and slow pathways and the impact of different slow recovery time constants (Fig. 11), it is clear that modest initial fractional entry into slow recovery pathways can still be associated with very robust use-dependent occupancy of slow recovery pathways.

#### Slow inactivation properties conferred by specific FGFs may differentially tune use-dependent changes in Nav availability

Given that each FGF isoform appears competent to interact with a variety of different SCNA subunits (Goldfarb, 2005; Goetz et al., 2009), any functional differences among FGF-A subunits or specificity in interactions with particular SCNA subunits are likely to generate profound differences on use-dependent changes in Nav availability. Based on the the simple calculations done here (Figs. 10,11), the two factors primarily affecting the extent and rate of development of use-dependent changes in Nav availability are, first, the initial differential rates of entry into fast and slow recovery pathways and, second, the rates of slow recovery from inactivation. Ultimately, careful understanding of such interactions of FGF-A isoforms with specific SCNA variants will be required. We propose that specific differences in the onset of inactivation and recovery from inactivation mediated by specific FGF-A isoforms may have profound affects on the extent of use-dependent accumulation of Nav channels in slow recovery from inactivation pathways. This, in turn, will differentially impact on the time course and extent of Nav availability during repetitive stimuli.

## Abbreviations

CCs: chromaffin cells
TTX: tetrodotoxin
AM: adrenal medulla
FGF: fibroblast growth factor homologous factor

## Acknowledgements

We thank the members of the Waxman group (Yale University) for providing Nav1.3 KO mice. We also thank Dr. Jeanne Nerbonne for making available FGF14 KO mice and very helpful discussions during the course of this work.

---

**Figure S1.**
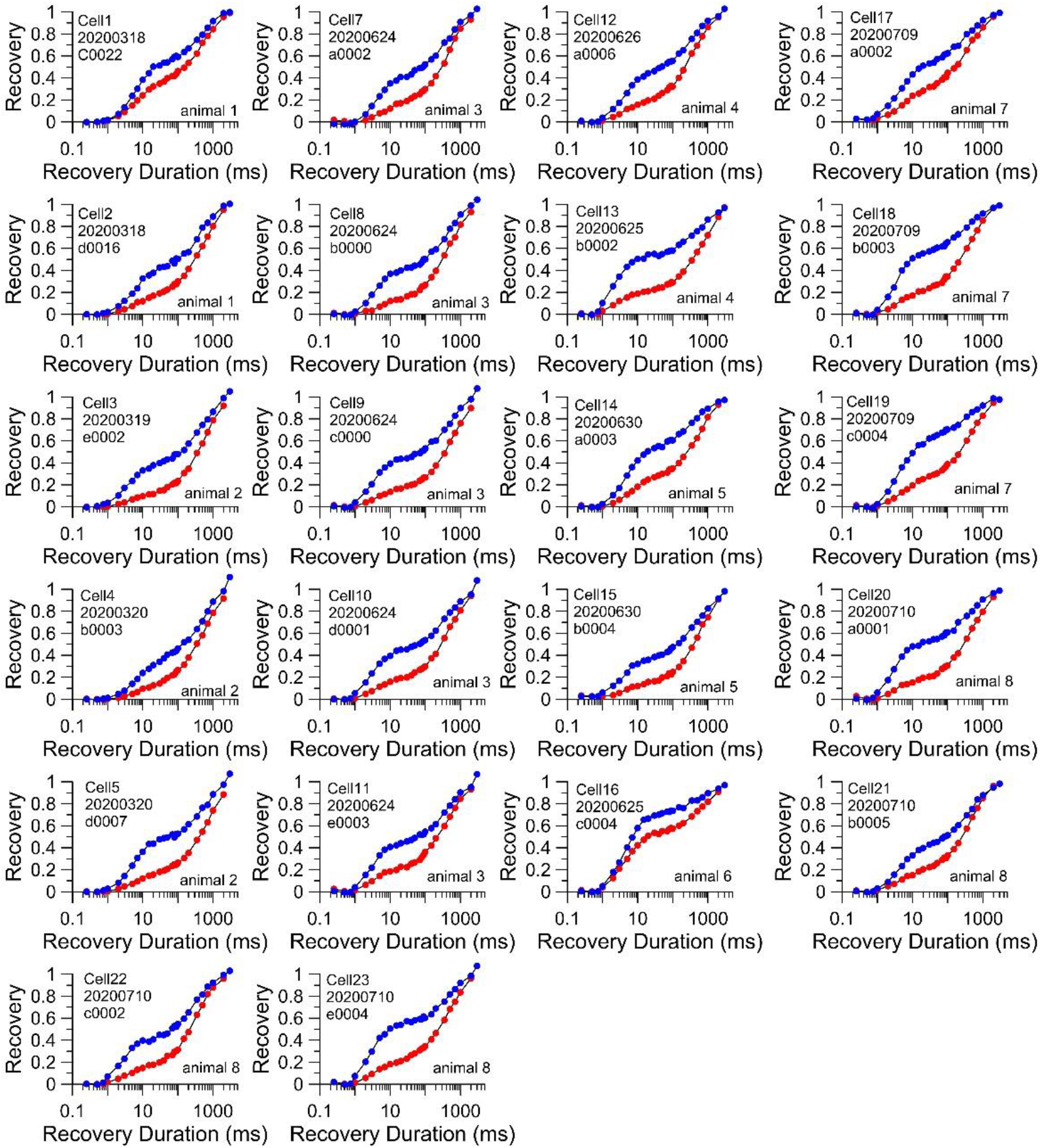
Examples of two-component recovery from inactivation following 1P and 10P test stimuli for WT cells. Each panel shows recovery from inactivation following a single 10 ms step to 0 mV (blue) and then following a 10 Hz train of 10 pulses to 0 mV (red). A total of 22 WT cells from 8 different animals were compared with the 1P and 10P protocols.

**Figure S2.**
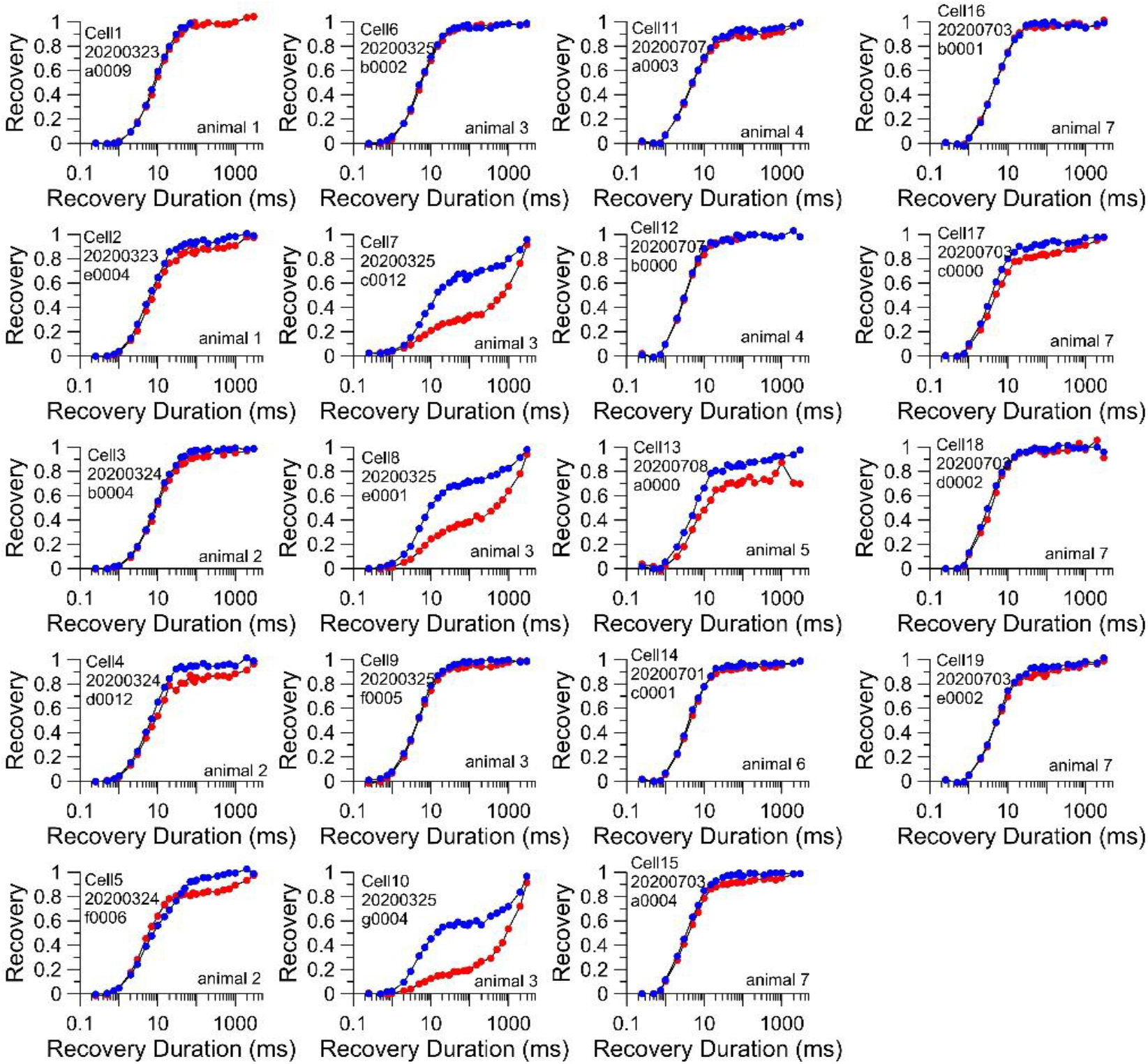
Examples of recovery from inactivation following 1P and 10P test stimuli for FGF14 KO cells. Each panel shows recovery from inactivation following a single 10 ms step to 0 mV (blue) and then following a 10 Hz train of 10 pulses to 0 mV (red). A total of 24 FGF14 KO cells from 7 different animals were compared with the 1P and 10P protocols. Three cells in column 2 exhibited clear two component recovery from inactivation following both 1P and 10P protocols. For these three cells, despite a smaller slow component of recovery following the 1P protocol, the 10P protocol resulted in a fraction of slow recovery comparable to or even larger than observed in WT cells (Fig. S1).

## References

Barbosa, C., and T.R. Cummins. 2016. Unusual Voltage-Gated Sodium Currents as Targets for Pain. Curr Top Membr. 78:599–638.

Bosch, M.K., Y. Carrasquillo, J.L. Ransdell, A. Kanakamedala, D.M. Ornitz, and J.M. Nerbonne. 2015. Intracellular FGF14 (iFGF14) Is Required for Spontaneous and Evoked Firing in Cerebellar Purkinje Neurons and for Motor Coordination and Balance. J Neurosci. 35:6752–6769.

Capes, D.L., M.P. Goldschen-Ohm, M. Arcisio-Miranda, F. Bezanilla, and B. Chanda. 2013. Domain IV voltage-sensor movement is both sufficient and rate limiting for fast inactivation in sodium channels. J Gen Physiol. 142:101–112.

Cummins, T.R., F. Aglieco, M. Renganathan, R.I. Herzog, S.D. Dib-Hajj, and S.G. Waxman. 2001. Nav1.3 sodium channels: rapid repriming and slow closed-state inactivation display quantitative differences after expression in a mammalian cell line and in spinal sensory neurons. Journal of Neuroscience. 21:5952–5961.

Cummins, T.R., S.D. Dib-Hajj, and S.G. Waxman. 2004. Electrophysiological properties of mutant Nav1.7 sodium channels in a painful inherited neuropathy. J Neurosci. 24:8232–8236.

Cummins, T.R., J.R. Howe, and S.G. Waxman. 1998. Slow closed-state inactivation: a novel mechanism underlying ramp currents in cells expressing the hNE/PN1 sodium channel. J Neurosci. 18:9607–9619.

Diwakar, S., J. Magistretti, M. Goldfarb, G. Naldi, and E. D’Angelo. 2009. Axonal Na^+^ channels ensure fast spike activation and back-propagation in cerebellar granule cells. J Neurophysiology. 101:519–532.

Dover, K., S. Solinas, E. D’Angelo, and M. Goldfarb. 2010. Long-term inactivation particle for voltage-gated sodium channels. Journal of Physiology. 588:3695–3711.

Eaholtz, G., T. Scheuer, and W.A. Catterall. 1994. Restoration of inactivation and block of open sodium channels by an inactivation gate peptide. Neuron. 12:1041–1048.

Effraim, P.R., J. Huang, A. Lampert, S. Stamboulian, P. Zhao, J.A. Black, S.D. Dib-Hajj, and S.G. Waxman. 2019. Fibroblast growth factor homologous factor 2 (FGF-13) associates with Nav1.7 in DRG neurons and alters its current properties in an isoform-dependent manner. Neurobiol Pain. 6:100029.

Fenwick, E.M., A. Marty, and E. Neher. 1982. Sodium and calcium channels in bovine chromaffin cells. J Physiol. 331:599–635.

Goetz, R., K. Dover, F. Laezza, N. Shtraizent, X. Huang, D. Tchetchik, A.V. Eliseenkova, C.F. Xu, T.A. Neubert, D.M. Ornitz, M. Goldfarb, and M. Mohammadi. 2009. Crystal structure of a fibroblast growth factor homologous factor (FHF) defines a conserved surface on FHFs for binding and modulation of voltage-gated sodium channels. J Biol Chem. 284:17883–17896.

Goldfarb, M. 2005. Fibroblast growth factor homologous factors: evolution, structure, and function. Cytokine Growth Factor Rev. 16:215–220.

Goldfarb, M. 2012. Voltage-gated sodium channel-associated proteins and alternative mechanisms of inactivation and block. Cell Mol Life Sci. 69:1067–1076.

Goldfarb, M., J. Schoorlemmer, A. Williams, S. Diwakar, Q. Wang, X. Huang, J. Giza, D. Tchetchik, K. Kelley, A. Vega, G. Matthews, P. Rossi, D.M. Ornitz, and E. D’Angelo. 2007. Fibroblast growth factor homologous factors control neuronal excitability through modulation of voltage-gated sodium channels. Neuron. 55:449–463.

Hamill, O.P., A. Marty, E. Neher, B. Sakmann, and F.J. Sigworth. 1981. Improved patch-clamp techniques for high-resolution current recording from cells and cell-free membrane patches. Pflugers Archiv. 391:85–100.

Hennessey, J.A., C.A. Marcou, C. Wang, E.Q. Wei, C. Wang, D.J. Tester, M. Torchio, F. Dagradi, L. Crotti, P.J. Schwartz, M.J. Ackerman, and G.S. Pitt. 2013. FGF12 is a candidate Brugada syndrome locus. Heart Rhythm. 10:1886–1894.

Herzog, R.I., T.R. Cummins, F. Ghassemi, S.D. Dib-Hajj, and S.G. Waxman. 2003. Distinct repriming and closed-state inactivation kinetics of Nav1.6 and Nav1.7 sodium channels in mouse spinal sensory neurons. J Physiol. 551:741–750.

Islas-Suarez, L., M. Gomez-Chavarin, R. Drucker-Colin, and A. Hernandez-Cruz. 1994. Properties of the sodium current in rat chromaffin cells exposed to nerve growth factor in vitro. J Neurophysiol. 72:1938–1948.

Klugbauer, N., L. Lacinova, V. Flockerzi, and F. Hofmann. 1995. Structure and functional expression of a new member of the tetrodotoxin-sensitive voltage-activated sodium channel family from human neuroendocrine cells. EMBO J. 14:1084–1090.

Laezza, F., B.R. Gerber, J.Y. Lou, M.A. Kozel, H. Hartman, A.M. Craig, D.M. Ornitz, and J.M. Nerbonne. 2007. The FGF14(F145S) mutation disrupts the interaction of FGF14 with voltage-gated Na+ channels and impairs neuronal excitability. J Neurosci. 27:12033–12044.

Laezza, F., A. Lampert, M.A. Kozel, B.R. Gerber, A.M. Rush, J.M. Nerbonne, S.G. Waxman, S.D. Dib-Hajj, and D.M. Ornitz. 2009. FGF14 N-terminal splice variants differentially modulate Nav1.2 and Nav1.6-encoded sodium channels. Mol Cell Neurosci. 42:90–101.

Lingle, C.J., P.L. Martinez-Espinosa, L. Guarina, and E. Carbone. 2017. Roles of Na+, Ca2+, and K+ channels in the generation of repetitive firing and rhythmic bursting in adrenal chromaffin cells. Pflugers Arch.

Liu, C.J., S.D. Dib-Hajj, M. Renganathan, T.R. Cummins, and S.G. Waxman. 2003. Modulation of the cardiac sodium channel Nav1.5 by fibroblast growth factor homologous factor 1B. J Biol Chem. 278:1029–1036.

Lou, X.L., X. Yu, X.K. Chen, K.L. Duan, L.M. He, A.L. Qu, T. Xu, and Z. Zhou. 2003. Na+ channel inactivation: a comparative study between pancreatic islet beta-cells and adrenal chromaffin cells in rat. J Physiol. 548:191–202.

Martinez-Espinosa, P., C. Yang, V. Gonzalez-Perez, X.M. Xia, and C.J. Lingle. 2014. Knockout of the BK beta2 subunit abolishes inactivation of BK currents in mouse adrenal chromaffin cells and results in slow-wave burst activity. Journal of General Physiology. 144:275–295.

Martinez-Espinosa, P.L., A. Neely, J.P. Ding, and C.J. Lingle. 2020. Fast inactivation of Na+ current in rat adrenal chromaffin cells involves two independent inactivation pathways. Journal of General Physiology. in preparation.

Milescu, L.S., T. Yamanishi, K. Ptak, and J.C. Smith. 2010. Kinetic properties and functional dynamics of sodium channels during repetitive spiking in a slow pacemaker neuron. Journal of Neuroscience. 30:12113–12127.

Munoz-Sanjuan, I., P.M. Smallwood, and J. Nathans. 2000. Isoform diversity among fibroblast growth factor homologous factors is generated by alternative promoter usage and differential splicing. J Biol Chem. 275:2589–2597.

Musa, H., C.F. Kline, A.C. Sturm, N. Murphy, S. Adelman, C. Wang, H. Yan, B.L. Johnson, T.A. Csepe, A. Kilic, R.S. Higgins, P.M. Janssen, V.V. Fedorov, R. Weiss, C. Salazar, T.J. Hund, G.S. Pitt, and P.J. Mohler. 2015. SCN5A variant that blocks fibroblast growth factor homologous factor regulation causes human arrhythmia. Proc Natl Acad Sci U S A. 112:12528–12533.

Navarro, M.A., A. Salari, J.L. Lin, L.M. Cowan, N.J. Penington, M. Milescu, and L.S. Milescu. 2020. Sodium channels implement a molecular leaky integrator that detects action potentials and regulates neuronal firing. Elife. 9.

Nemoto, T., T. Yanagita, T. Maruta, C. Sugita, S. Satoh, T. Kanai, A. Wada, and M. Murakami. 2013. Endothelin-1-induced down-regulation of NaV1.7 expression in adrenal chromaffin cells: attenuation of catecholamine secretion and tau dephosphorylation. FEBS Lett. 587:898–905.

Pablo, J.L., and G.S. Pitt. 2014. Fibroblast Growth Factor Homologous Factors: New Roles in Neuronal Health and Disease. Neuroscientist.

Pablo, J.L., C. Wang, M.M. Presby, and G.S. Pitt. 2016. Polarized localization of voltage-gated Na+ channels is regulated by concerted FGF13 and FGF14 action. Proc Natl Acad Sci U S A.

Rush, A.M., E.K. Wittmack, L. Tyrrell, J.A. Black, S.D. Dib-Hajj, and S.G. Waxman. 2006. Differential modulation of sodium channel Na(v)1.6 by two members of the fibroblast growth factor homologous factor 2 subfamily. Eur J Neurosci. 23:2551–2562.

Siekierska, A., M. Isrie, Y. Liu, C. Scheldeman, N. Vanthillo, L. Lagae, P.A. de Witte, H. Van Esch, M. Goldfarb, and G.M. Buyse. 2016. Gain-of-function FHF1 mutation causes early-onset epileptic encephalopathy with cerebellar atrophy. Neurology. 86:2162–2170.

Smallwood, P.M., I. Munoz-Sanjuan, P. Tong, J.P. Macke, S.H. Hendry, D.J. Gilbert, N.G. Copeland, N.A. Jenkins, and J. Nathans. 1996. Fibroblast growth factor (FGF) homologous factors: new members of the FGF family implicated in nervous system development. Proc Natl Acad Sci U S A. 93:9850–9857.

Solaro, C.R., M. Prakriya, J.P. Ding, and C.J. Lingle. 1995. Inactivating and noninactivating Ca^2+^- and voltage-dependent K^+^ current in rat adrenal chromaffin cells. Journal of Neuroscience. 15:6110–6123.

Tamura, R., T. Nemoto, T. Maruta, S. Onizuka, T. Yanagita, A. Wada, M. Murakami, and I. Tsuneyoshi. 2014. Up-regulation of NaV1.7 sodium channels expression by tumor necrosis factor-alpha in cultured bovine adrenal chromaffin cells and rat dorsal root ganglion neurons. Anesth Analg. 118:318–324.

Vandael, D.H., M.M. Ottaviani, C. Legros, C. Lefort, N.C. Guerineau, A. Allio, V. Carabelli, and E. Carbone. 2015. Reduced availability of voltage-gated sodium channels by depolarization or blockade by tetrodotoxin boosts burst firing and catecholamine release in mouse chromaffin cells. J Physiol. 593:905–927.

Venkatesan, K., Y. Liu, and M. Goldfarb. 2014. Fast-onset long-term open-state block of sodium channels by A-type FHFs mediates classical spike accommodation in hippocampal pyramidal neurons. J Neurosci. 34:16126–16139.

Wada, A., E. Wanke, F. Gullo, and E. Schiavon. 2008. Voltage-dependent Na(v)1.7 sodium channels: multiple roles in adrenal chromaffin cells and peripheral nervous system. Acta Physiol (Oxf). 192:221–231.

Wada, A., T. Yanagita, H. Yokoo, and H. Kobayashi. 2004. Regulation of cell surface expression of voltage-dependent Nav1.7 sodium channels: mRNA stability and posttranscriptional control in adrenal chromaffin cells. Front Biosci. 9:1954–1966.

Wang, C., B.C. Chung, H. Yan, S.Y. Lee, and G.S. Pitt. 2012. Crystal structure of the ternary complex of a NaV C-terminal domain, a fibroblast growth factor homologous factor, and calmodulin. Structure. 20:1167–1176.

Wang, Q., M.E. Bardgett, M. Wong, D.F. Wozniak, J. Lou, B.D. McNeil, C. Chen, A. Nardi, D.C. Reid, K. Yamada, and D.M. Ornitz. 2002. Ataxia and paroxysmal dyskinesia in mice lacking axonally transported FGF14. Neuron. 35:25–38.

Wang, X., H. Tang, E.Q. Wei, Z. Wang, J. Yang, R. Yang, S. Wang, Y. Zhang, G.S. Pitt, H. Zhang, and C. Wang. 2017. Conditional knockout of Fgf13 in murine hearts increases arrhythmia susceptibility and reveals novel ion channel modulatory roles. J Mol Cell Cardiol.

West, J.W., D.E. Patton, T. Scheuer, Y. Wang, A.L. Goldin, and W.A. Catterall. 1992. A cluster of hydrophobic amino acid residues required for fast Na(+)-channel inactivation. Proc Natl Acad Sci U S A. 89:10910–10914.

Xiao, M., M.K. Bosch, J.M. Nerbonne, and D.M. Ornitz. 2013. FGF14 localization and organization of the axon initial segment. Mol Cell Neurosci. 56:393–403.

Yan, Z., Q. Zhou, L. Wang, J. Wu, Y. Zhao, G. Huang, W. Peng, H. Shen, J. Lei, and N. Yan. 2017. Structure of the Nav1.4-beta1 Complex from Electric Eel. Cell. 170:470–482 e411.

Yanagita, T., T. Maruta, Y. Uezono, S. Satoh, N. Yoshikawa, T. Nemoto, H. Kobayashi, and A. Wada. 2007. Lithium inhibits function of voltage-dependent sodium channels and catecholamine secretion independent of glycogen synthase kinase-3 in adrenal chromaffin cells. Neuropharmacology. 53:881–889.

Yang, C.T., X.H. Zeng, X.M. Xia, and C.J. Lingle. 2009. Interactions between beta subunits of the KCNMB family and Slo3: beta4 selectively modulates Slo3 expression and function. PLoS One. 4:e6135.

Yang, J., Z. Wang, D.S. Sinden, X. Wang, B. Shan, X. Yu, H. Zhang, G.S. Pitt, and C. Wang. 2016. FGF13 modulates the gating properties of the cardiac sodium channel Nav1.5 in an isoform-specific manner. Channels (Austin). 10:410–420.

Zhang, Q., M.V. Chibalina, M. Bengtsson, L.N. Groschner, R. Ramracheya, N.J. Rorsman, V. Leiss, M.A. Nassar, A. Welling, F.M. Gribble, F. Reimann, F. Hofmann, J.N. Wood, F.M. Ashcroft, and P. Rorsman. 2014. Na+ current properties in islet alpha- and beta-cells reflect cell-specific Scn3a and Scn9a expression. Journal of Physiology (Lond). 592:4677–4696.

